# *PTCH1* mutant human cerebellar organoids are associated with altered neural development and early pathways of medulloblastoma oncogenesis

**DOI:** 10.1101/2023.05.10.540200

**Authors:** Max J. van Essen, Joey Riepsaame, Sally A. Cowley, John Jacob, Esther B. E. Becker

**Author notes:** Co-senior author (John Jacob, Esther B. E. Becker).

## Abstract

Patched 1 (PTCH1) is the primary receptor for Sonic Hedgehog (SHH) ligand and negatively regulates SHH signalling, an essential pathway in human embryogenesis. Loss-of-function mutations in *PTCH1* are associated with altered neuronal development and the malignant brain tumour medulloblastoma (MB). As a result of differences between murine and human development, molecular and cellular perturbations that arise from human *PTCH1* mutations remain poorly understood. Here, we employ cerebellar organoids differentiated from human induced pluripotent stem cells (iPSC) combined with CRISPR/Cas9 gene editing to investigate the earliest molecular and cellular consequences of *PTCH1* mutations on human cerebellar development. Our findings support the occurrence of developmental mechanisms in cerebellar organoids that mirror *in vivo* processes of regionalisation and SHH signalling, and offer new insight into early pathophysiological events of MB tumorigenesis.

**Higlights:**

- Differentiation of human iPSC into cerebellar organoids
- Homozygous LOF of *PTCH1* prevents cerebellar organoid differentiation
- *PTCH1+/-* cerebellar organoids display tissue-specific effects of SHH signalling
- Early altered gene expression relevant for MB in *PTCH1+/-* cerebellar organoids

## Introduction

Patched 1 (PTCH1) is the primary receptor for Sonic Hedgehog (SHH) ligand and the most proximal negative regulator of the SHH signal transduction pathway. SHH signalling is essential in human embryogenesis and regulates differentiation and proliferation of target cells (Fuccillo et al., 2006). During cerebellar development, SHH activity regulates the proliferative expansion of cerebellar granule cell progenitors (GCP) in the external granular layer (EGL) of the cerebellum (Aguilar et al., 2012; Dahmane and Ruiz i Altaba, 1999; Leto et al., 2016; Wechsler-Reya and Scott, 1999). PTCH1 plays a pivotal role in controlling SHH-induced proliferation by preventing Smoothened (SMO) from transducing SHH signals (Blassberg and Jacob, 2017; Corbit et al., 2005; Corcoran and Scott, 2006; Denef et al., 2000; Dwyer et al., 2007; Khaliullina et al., 2009; Nachtergaele et al., 2012). Consequently, PTCH1 loss-of-function (LOF) leads to constitutive SMO activation and SHH signalling. SHH signalling primarily occurs through post-trascriptional modulation of GLI proteins, a class of transcription factors that regulate stem cell maintenance, proliferation, and differentiation (Ruiz i Altaba et al., 2002a).

*PTCH1* LOF gene mutations are associated with various types of cancer (Ruiz i Altaba et al., 2002b) including the cerebellar brain tumour medulloblastoma (MB), the most frequent malignant brain tumour in children (Ostrom et al., 2018; Smoll and Drummond, 2012). A major MB subtype arises from overactive SHH signalling in GCP that is classified as SHH-MB. Notably, around 44% of all SHH-MB tumours harbour heterozygous LOF mutations in *PTCH1* (Garcia-Lopez et al., 2021; Northcott et al., 2019). Mouse and human studies have helped to identify GCP as the cell-of-origin of SHH-MB (Hovestadt et al., 2019; Vladoiu et al., 2019; Yang et al., 2008), but early molecular and cellular perturbations resulting from human *PTCH1* mutations that drive later malignant transformation remain incompletely understood. Increasing evidence reveals how genetic events have different effects in mice compared to humans, which emphasises the importance of developing tools to study human development and disease (Bouaoun et al., 2016; Jacks et al., 1994). Furthermore, studies in recent years have shed light on relevant differences between murine and human cerebellar and MB development. Specifically, Purkinje cells (PC) in mice secrete Shh that acts on GCP to induce proliferation (Dahmane and Ruiz i Altaba, 1999), while human PC are likely not mature enough to produce SHH when SHH-induced proliferation first occurs (Zecevic and Rakic, 1976). Thus, human-specific models are needed to better understand PTCH1-mediated signalling relevant to human disease and to make further advances in MB treatment.

Induced pluripotent stem cells (iPSC) technology has transformed the way human development and disease is studied and opened new human-centric avenues for disease modelling. Unlike transformed cell lines, iPSC allow the *in vitro* study of human cellular differentiation and physiology under conditions of a normal karyotype, gene dosage, cell cycle and metabolic profile. Among the numerous cell types and tissues that can now be generated *in vitro* are cerebellar neurons and organoids that recapitulate early hindbrain patterning and cerebellar lineage formation (Muguruma et al., 2015; Watson et al., 2018). Single-cell RNA sequencing of cerebellar organoids maintained for 90 days *in vitro* (DIV) revealed the presence of all major cerebellar developmental neuronal cell types including rhombic lip (RL), GCP, and proliferating and post-mitotic granule cells (GCs) (Nayler et al., 2021). By combining cerebellar organoid technology with CRISPR/Cas9 gene editing, we aimed to investigate the earliest molecular and cellular consequences of *PTCH1* mutations on cerebellar development by cerebellar organoid differentiation.

CRISPR gene editing has emerged as an exciting tool for functional gene studies and disease modelling (van Essen et al., 2021). Using CRISPR technology, mutant iPSC lines can be generated and compared to isogenic healthy controls, which avoids the confounding effect of different genetic backgrounds on the mutant cellular phenotype. Although groups have modelled SHH-MB using neuroepithelial stem cells (NES) differentiated from iPSC from patients carrying *PTCH1* mutations (Huang et al., 2019; Susanto et al., 2020), these studies required orthotopic injection in mice. Directing *PTCH1* heterozygous and homozygous mutant iPSC to cerebellar organoid differentiation allows the investigation of *PTCH1* dysfunction on cerebellar development for the first time in a human model *in vitro*. The present study describes the generation of *PTCH1* mutant iPSC using CRISPR targeting, and investigates the effects of *PTCH1* LOF on cerebellar differentiation. We find that homozygous *PTCH1* LOF in iPSC prevents cerebellar differentiation and promotes ventral forebrain identity through early high-level SHH signalling. In contrast, *PTCH1* heterozygous iPSC differentiate into cerebellar organoids that harbour an expanded RL and GCP population and display features associated with pre-neoplastic stages of MB. Together, these results illustrate the utility of cerebellar organoids in studying the effects of human gene mutations on development and disease.

## Results

### Introduction of a *PTCH1* LOF mutation in human induced pluripotent stem cells

To generate *PTCH1* mutant human iPSC, we targeted exon 3 of *PTCH1* (a gene with 24 exons), which encodes an essential region of the protein. Using a pair of gRNAs and a Cas9 ribonucleoprotein (RNP) approach, we excised the splice donor (SD) of this exon and generated *PTCH1* mutant iPSC (Figure 1A). We established monoclonal heterozygous and homozygous iPSC lines by single-cell cloning on mouse embryonic feeder (MEF) cells. Each line was genotyped using PCR to detect cutting at the gRNA target sites (Figure 1B), which was confirmed by Sanger sequencing (Figure S1). Examination of cDNA generated through reverse translation of *PTCH1* mRNA from mutant human iPSC clones revealed that CRISPR/Cas9 induced alternative splicing and exclusion of exon 3 (Figure 1C). As exon 3 is an asymmetric exon (not a multiple of three base pairs), this results in a -1 frameshift in the mutant mRNA with multiple premature stop-codons in the new reading frame and thus a *PTCH1* null mutant.

**Figure 1.**
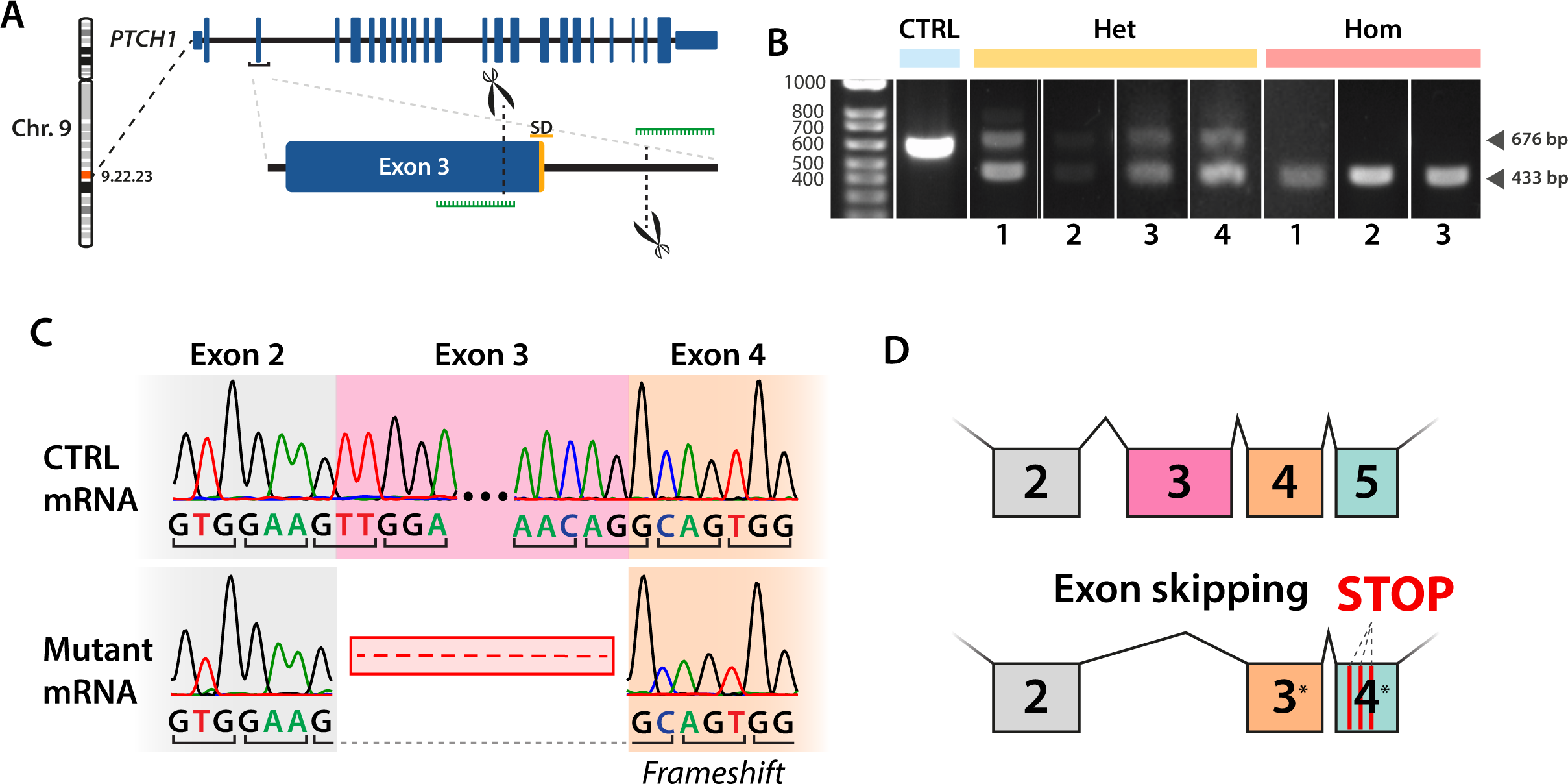
Generation of *PTCH1* loss-of-function mutations in human induced pluripotent stem cells using CRISPR (A) CRISPR gene editing strategy to target exon 3 of the *PTCH1* gene. Guide RNAs (green) were designed to result in the excision of a 243-bp region spanning the splice donor site (SD) of exon 3. (B) PCR genotyping confirmed the generation of heterozygous (Het) *PTCH1* iPSC clones producing both control (CTRL) (676bp) and mutant (433bp) PCR amplicons as well as homozygous (Hom) clones producing only the mutant amplicon. (C) Sanger sequencing of *PTCH1* mRNA revealed the exclusion of exon 3 in the mutant mRNA resulting in a frameshift mutation in exon 4. (D) Schematic depiction of the effect of exon 3 exclusion, resulting in a frameshift mutation and multiple premature translation termination codons in the new exon 4. See also Figures S1-S5.

*PTCH1* mutant clones maintained iPSC morphology and expressed NANOG and Tra-1-60 to similar levels, as measured by flow cytometry (Figure S2). A screen for chromosomal aberrations using a SNP-array testing over 600,000 sites in the human genome did not reveal any new amplifications, deletions, or re-arrangements (Figure S3). Furthermore, no specific off-target effects of the CRISPR strategy were detected upon sequencing of the top five off-target sites for each of the gRNAs (Figure S4).

PTCH1 LOF is known to increase SHH signalling in SHH-responsive cells (Chen et al., 2002; Goodrich et al., 1997; Taipale et al., 2000). Although iPSC are not known to be SHH responsive, we used reverse transcriptase quantitative PCR (RT-qPCR) to measure mRNA levels of well-established readouts of SHH pathway activity, *GLI1* and *PTCH1* (Goodrich et al., 1996; Lee et al., 1997). Both targets were modestly upregulated in homozygous (*PTCH1*-/-) iPSC compared to isogenic control cells (*GLI1* p<0.001; *PTCH1* p=0.01; Figure S5A) consistent with the absence of PTCH1-SHH signalling. In heterozygous (*PTCH1*+/-) iPSC, only *GLI1* expression was elevated in comparison with control iPSC (p=0.027). To test whether these upregulated readouts of SHH signalling implied SHH induced proliferation of our iPSC lines, we analysed the proliferation of *PTCH1* mutant iPSC, by exposing cells to a pulse of 5-ethynyl-2’-deoxyuridine (EdU). No statistical differences could be observed in the proportions of cells in different stages of the cell cycle (Figure S5B). Together, these results show that *PTCH1* mutant iPSC maintain normal morphology and proliferation.

### Cerebellar differentiation of *PTCH1* mutant iPSC reveals changes in morphology and cerebellar gene expression

To investigate the effect of PTCH1 LOF on cerebellar differentiation, control and mutant *PTCH1* human iPSC lines were differentiated into cerebellar organoids following a previously published protocol by our laboratory (Nayler et al., 2021) (Figure 2A). Heterozygous organoids exhibited a similar shape to control organoids (Figure 2B). In contrast, homozygous organoids showed changed morphology with more irregular growth and polarisation of the tissue, marked by the greater translucency of the edges on brightfield imaging. Comparing sizes of mutant versus control organoids respectively, we found that *PTCH1* heterozygous and homozygous mutants grew more quickly compared to control organoids (Figure 2C). These results are in line with previous studies that show neural overgrowth in *Ptch1*+/- mice (Jackson et al., 2020; Oliver et al., 2005). The marked altered morphology of *PTCH1*-/- organoids suggested changes in the differentiation trajectory of the homozygous mutant iPSC.

**Figure 2.**
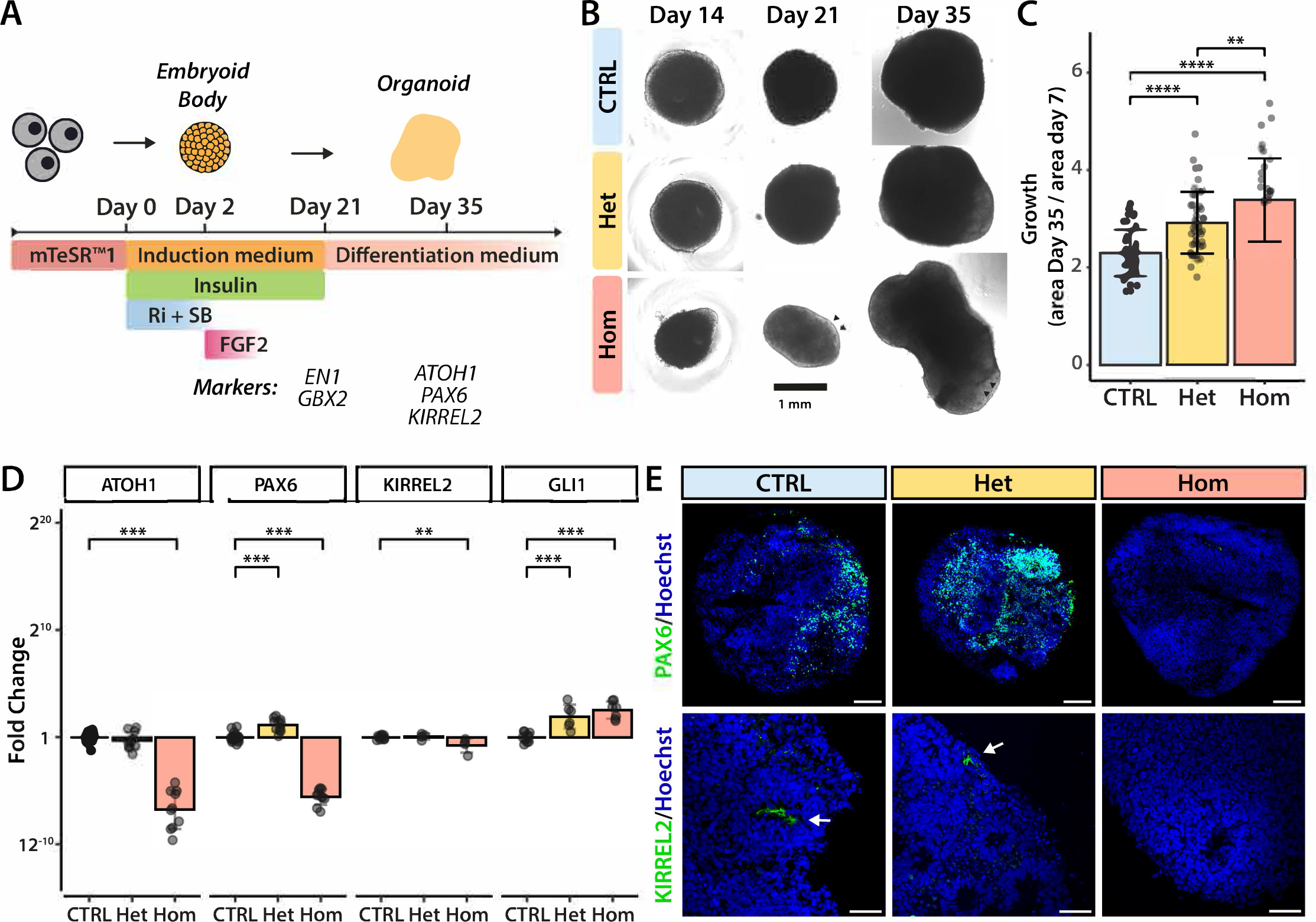
Cerebellar differentiation of *PTCH1* mutant iPSC reveals changes in morphology and cerebellar lineage markers (A) Schematic overview of the cerebellar differentiation protocol. (B) Representative images of organoids generated from control (CTRL), *PTCH1+/-* (Het) and *PTCH1-/-* (Hom) iPSC during differentiation. Arrowheads indicate regions of enhanced polarisation within the homozygous organoids. (C) Relative size change of organoids at day 35 compared to day 7. Data are composed of 46 (CTRL), 48 (Het) and 35 (Hom) individual organoids across three differentiations. Statistical significance is indicated as *p<0.05, **p<0.01, ***p<0.001, computed by ANOVA with Tukey post-hoc test. Error bars indicate standard deviation. (D) Expression of cerebellar and SHH-pathway genes, measured by RT-qPCR. Each data point represents an experimental replicate. Data are composed of at least three biological replicates from separate differentiations. *p<0.05, **p<0.01, ***p<0.001, computed with ANOVA followed by Dunnett’s post-hoc test.. Error bars indicate standard deviation. E) Immunofluorescence staining of Day 35 organoids with antibodies specific to the rhombic lip lineage marker PAX6 (green) and ventricular zone marker KIRREL2 (green). KIRREL2 is primarily expressed in the small neuroepithelial lumina present in cerebellar organoids. Nuclei are visualised in blue by Hoechst staining. Scale bars 100μm in the top row images and 50μm in the bottom row images. See also Figure S2.

We next assessed differentiation along the midbrain-hindbrain lineage by measuring mRNA levels of Engrailed 1 (EN1), which marks cells with midbrain and anterior hindbrain region identity, and gastrulation brain homeobox 2 (GBX2), which is expressed by anterior hindbrain cells (Liu and Joyner, 2001). Expression of Orthodenticle homeobox 2 (OTX2), which is expressed in the developing midbrain but absent in the presumptive cerebellar territory was also used to distinguish the two regions (Vernay et al., 2005). At day 21 of the differentiation protocol, the expression of *EN1* and *GBX2* was increased in organoids of all three genotypes compared to levels in iPSC, while *OTX2* levels remained low. Between organoids of the different genotypes, only expression of *EN1* was significantly higher in *PTCH1*+/*-* and *PTCH1*-/- organoids compared to controls (p=0.01 for both genotypes) (Figure S6). To further explore the impact of *PTCH1* LOF on cerebellar induction we analysed the expression of cerebellar lineage markers after 35 days of differentiation. Gene expression of *ATOH1* and *PAX6*, which mark the glutamatergic lineage, and *KIRREL2* expressed by presumptive GABAergic neurons were measured by RT-qPCR (Figure 2D). Interestingly, we found that expression of both glutamatergic and GABAergic cerebellar markers was significantly lower in homozygous organoids compared to control. Loss of PAX6 and KIRREL2 expression in homozygous organoids was confirmed by immunofluorescence (Figure 2E). In contrast, heterozygous organoids expressed *ATOH1* and *KIRREL2* mRNA at similar levels to controls. In fact, the expression of *PAX6* mRNA was higher in heterozygous, compared to control organoids (p<0.001), opposite to the effect seen in homozygous organoids. These changes were accompanied by increased SHH-pathway activity in both heterozygous and homozygous genotypes as measured by the upregulated expression of *GLI1* (Figure 2D). The lower expression of cerebellar lineage markers in *PTCH1*-/- organoids suggests that cell patterning is severely disrupted. By contrast, the maintained expression of cerebellar markers in *PTCH1*+/- heterozygous organoids implies that sufficient PTCH1 activity is present in these organoids to facilitate the initial stages of cerebellar development.

### Homozygous *PTCH1* loss-of-function prevents cerebellar organoid differentiation

The observed changes in cerebellar markers, organoid growth rate, and morphology indicate significant changes in organoid development as a result of PTCH1 loss. To investigate these further, 35-day-old control and *PTCH1* mutant organoids were processed for bulk RNA sequencing. Using principal component analysis (PCA), we investigated transcriptome similarity and determined that samples clustered by genotype, with heterozygous samples locating closer to controls than to homozygous organoids (Figure 3A). When we compared gene expression of control organoids with homozygous mutants, 4579 genes met the adjusted p-value cut-off (<0.05) of differential expression (Data 1). Interestingly, genes associated with the ventral neural tube (*SHH*, *FOXA1* and *FOXA2*) were among the most upregulated genes in homozygous organoids, while genes marking the dorsal neural tube (*WNT3A* and *TLX3*) were among the most downregulated (Figure 3B). Further analysis revealed ventral markers were highly expressed in homozygous organoids while the expression of dorsal genes was lacking, compared to control organoids which showed an opposite pattern (Figure 3C). Using immunofluorescence staining, we confirmed the ectopic expression of the ventral neural tube marker NKX2-2 in homozygous *PTCH1*-/- mutant organoids but not in controls (Figure 3D,E). These regions are suggestive of ventral patterning in homozygous organoids similar to that seen in the developing neural tube (Briscoe and Ericson, 1999). When we analysed the expression of genes associated with regionalisation of the neural tube, *PTCH1*-/- organoids displayed upregulation of genes related to the forebrain, including *NKX2-1, FOXG1* and *SIX3*, and downregulation of hindbrain markers (for example *GBX2, ATOH1,* and *LMX1A)* compared to controls (Figure 3C). Consistent with a change in neural differentiation trajectory, increased expression of WNT antagonist *DKK1* and downregulation of WNT readouts *AXIN2* and *LGR5* was observed, a process that is critical for the formation of forebrain structures (del Barco Barrantes et al., 2003; Glinka et al., 1998). Together, these findings indicate that absence of PTCH1 function results in striking ventralisation and an anterior shift from mid-hindbrain to forebrain identity.

**Figure 3.**
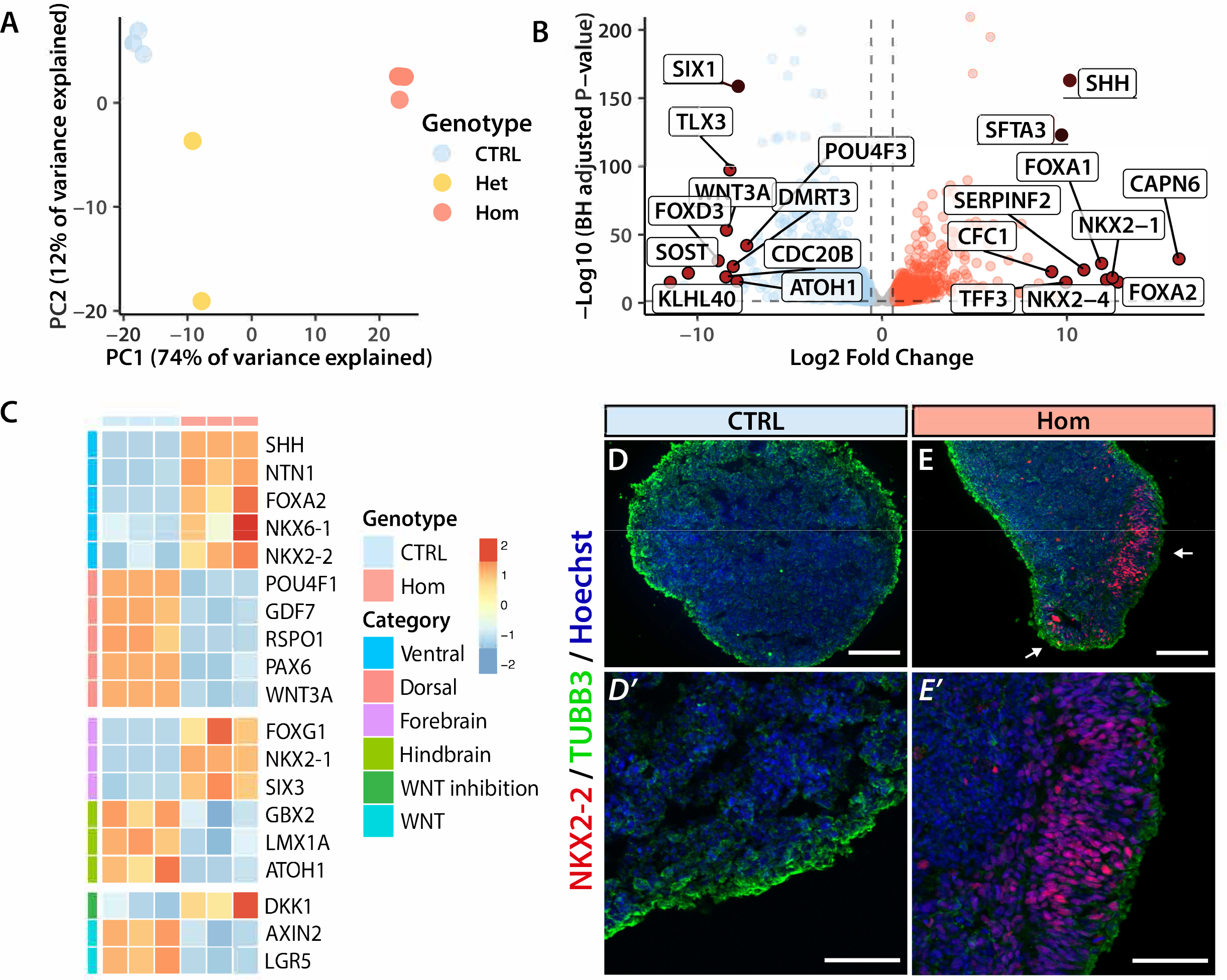
Homozygous *PTCH1* loss-of-function prevents cerebellar organoid differentiation (A) Principal component analysis using the 1000 most variable genes showing the separation of samples in principal component (PC) 1 and PC2. Control (CTRL) samples were generated with the AH017 line, *PTCH1+/-* (Het) samples with lines A3 and C6, and *PTCH1-/-* (Hom) samples with lines A2, B3, and H3. (B) Volcano plot of the ten genes with the most significant change in either direction. Genes upregulated in homozygous organoids are depicted in red, downregulated genes are blue. (C) Expression of genes associated with the ventral neural tube (Ventral), dorsal neural tube (Dorsal), hindbrain, forebrain, WNT inhibition and WNT signalling. Values are scaled within rows. The heatmap shows CTRL samples generated with the AH017 line, and Hom (*PTCH1-/-*) samples generated with clones A2, B3, and H3. (D-E) Immunofluorescence staining of day 35 organoids with antibodies specific to pan-neuronal marker TUBB3 (green) and ventral neural tube marker NKX2-2 (red). Nuclei are visualised in blue by Hoechst staining. Scale bars in D and E are 250μm, and in D’ and E’ 100μm. White arrows marks regions of NKX2-2 expression.

### The altered differentiation trajectory in *PTCH1***-/-** organoids is SHH signalling-dependent

The role of SHH signalling in ventralisation of the neural tube has been described extensively (Patten and Placzek, 2000) and suggests the altered differentiation trajectory in *PTCH1*-/- organoids is likely caused by activated SHH signalling. We conducted experiments to either activate or repress SHH signalling in control and mutant organoids, with the aim of clarifying the role of SHH in cerebellar patterning early in development. First, we exposed control and homozygous organoids to a SHH signalling inhibitor and determined whether the ventralisation of *PTCH1*-/- organoids could be blocked. During the first 35 days of differentiation, control and homozygous organoids were exposed to either DMSO or Cyclopamine, which inhibits SHH signal transduction downstream of PTCH1 and at the level of SMO (32) (Figure 4A,B). Cyclopamine treatment had only minor effects on gene expression in control organoids. In *PTCH1*-/- organoids, Cyclopamine rescued the expression of both *ATOH1* and *PAX6* in a dose-dependent manner, while moderately reducing *SHH* expression (Figure 4C). Protein expression of PAX6 and NKX2-2, which *in vivo* are used as markers of dorsal and ventral neural tube specification, respectively was determined using immunofluorescence staining (Figure 4D). This showed a marked increase in expression of PAX6 in homozygous organoids, surpassing PAX6 expression in control organoids. NKX2-2 expression was eliminated, and KIRREL2 immunostaining was rescued, whereas expression of TUBB3, a pan-neuronal marker, remained unchanged (Figure 4D). These results confirm the change of differentiation trajectory in *PTCH1*-/- organoids to be SHH-dependent and demonstrate how expression of cerebellar markers can be rescued in homozygous organoids by antagonizing SHH signalling.

**Figure 4.**
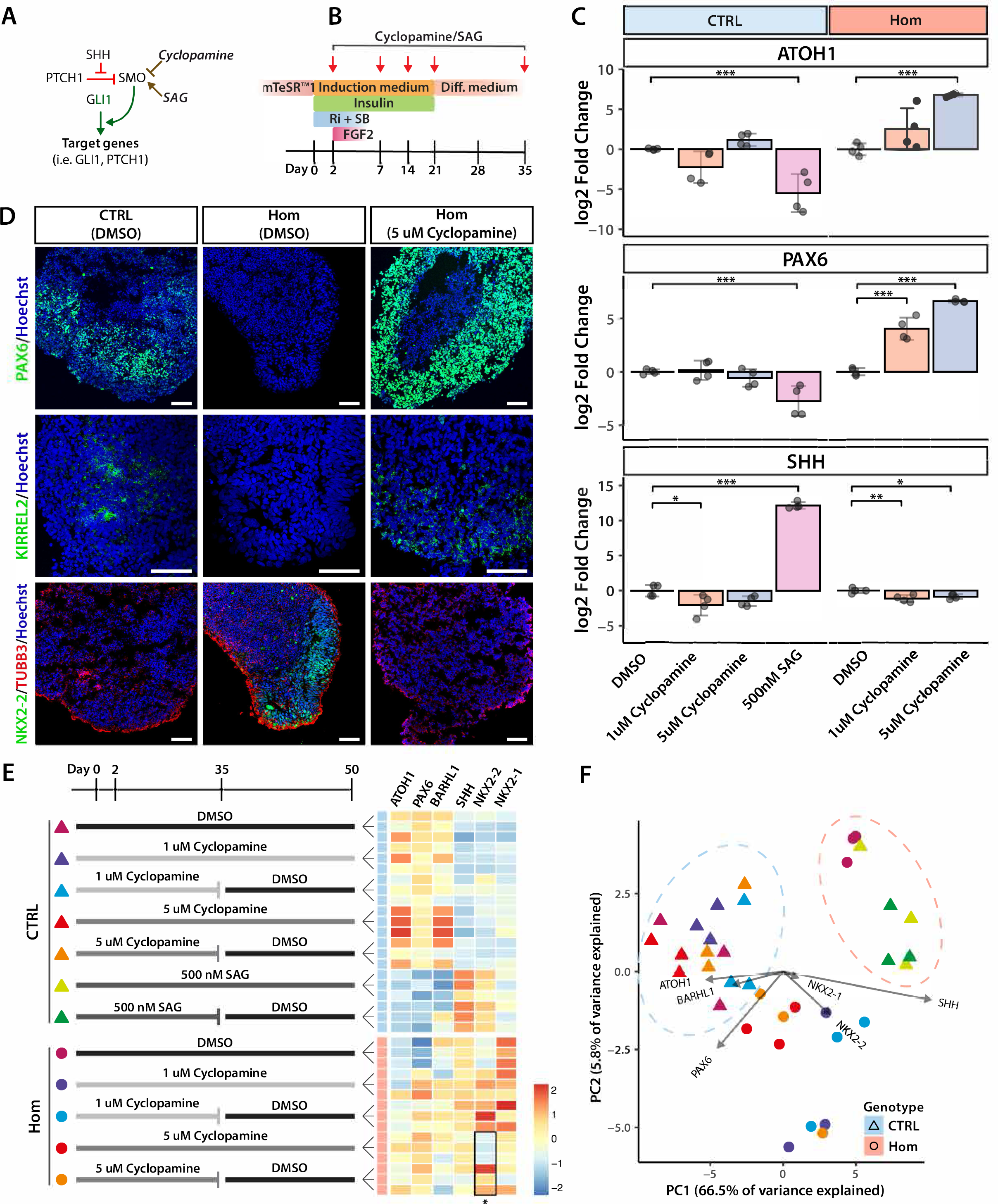
**SHH pathway inhibition rescues cerebellar marker expression** (A) Simplified schematic of the SHH pathway with the effect of Cyclopamine (red line, inhibitory) and SAG (green arrow, activating) on SMO indicated. (B) Schematic of experimental design to test the effect of SHH pathway inhibition on *PTCH1*-/- cerebellar organoid differentiation. (C) Gene expression of day 35 organoids in different treatment conditions as measured by RT-qPCR. Expression is relative to the DMSO control condition of each line and normalized using *ACTB* and *GAPDH*. Statistical significance was determined using ANOVA followed by Dunnett’s post-hoc test. *p<0.05, **p<0.01, ***p<0.001. n=4 per condition from two differentiations. Error bars indicate standard deviation. (D) Immunofluorescence staining of day 35 organoids from different conditions with antibodies specific to PAX6 (green), KIRREL2 (green) or TUBB3 (red) and NKX2-2 (green). Nuclei are visualised in blue by Hoechst staining. Scale bars 100μm. (E) Experimental design to test the effect of Cyclopamine or SAG treatment cessation on cerebellar marker expression is depicted on the left. Heatmap showing the ΔCt values normalised to ACTB and GAPDH in each condition. Asterisk indicates significance (p<0.05) computed by a two-tailed t-test (n=3) in comparing treatment cessation with continued treatment. Values are scaled within columns. (F) Principal component analysis showing different treatment and genotype conditions. The contribution of each gene to the principal components PC1 and PC2 is indicated by an arrow. The length of the arrow relates to the size of the contribution, indicating *SHH* expression contributes most to changes in PC1 and *PAX6* expression having the greatest effect on PC2.

Our data are consistent with the hypothesis that early, high-level SHH signalling results in a cell-fate switch to a ventral forebrain-like identity marked by the expression of *NKX2-2* and *NKX2-1* that could be reversed by treatment with Cyclopamine. This raises the question as to whether Cyclopamine treatment of homozygous organoids could permanently rescue cerebellar differentiation. We therefore investigated the effects on gene expression of relief from SHH inhibition upon the continued culture of Cyclopamine-treated organoids for another 15 days in the absence of the Smo inhibitor (Figure 4E). We found that expression of cerebellar GC markers *ATOH1*, *PAX6,* and *BARHL1* at day 50 did not significantly decrease in organoids when Cyclopamine treatment was discontinued after 35 days. Furthermore, *SHH* and *NKX2- 1* remained at similar levels in organoids exposed to either the 35-day or 50-day Cyclopamine treatment regimen. Most notably, *NKX2-*2 expression re-emerged after drug washout in homozygous organoids treated with 5μM Cyclopamine for 35 days. In other experiments, control organoids were treated with 500nM SAG to simulate enhanced SHH stimulation. SAG treatment significantly increased *SHH* expression and decreased mRNA levels of *ATOH1* and *PAX6*, similar to the changes seen in homozygous *PTCH1*-/- organoids (Figure 4C). These changes in gene expression persisted to at least day 50 even when SAG treatment was withdrawn after day 35.

We also performed PCA of the ΔCT values generated by RT-qPCR to examine the transcriptional changes comprehensively. Control organoids exposed to SAG treatment projected towards *PTCH1*-/- DMSO-treated organoids, suggesting similarity in gene expression (Figure 4F). Cyclopamine treatment caused homozygous mutant organoids to cluster away from DMSO-treated homozygous organoids in PC1, depending on the dose of Cyclopamine. Thus, SHH signalling appears to be the main driver of the gene expression profile differences between control and *PTCH1*-/- homozygous organoids. This is corroborated by *SHH* being the main contributing gene in PC1, while cerebellar GCP genes *ATOH1*, *PAX6*, and *BARHL1* all act in the opposing direction. Taken together, these results show that high-level and early SHH signalling induces a ventral-forebrain differentiation trajectory in *PTCH1*-/- homozygous organoids. This effect can be prevented by pharmacological SHH inhibition resulting in lasting rescue of the dorsal hindbrain phenotype associated with cerebellar differentiation.

### *PTCH1*+/- cerebellar organoids display tissue-specific effects of increased SHH signalling

Next, we investigated transcriptomic changes in day 35 heterozygous *PTCH1*+/- organoids. Because of sample dropout, we performed an additional round of sequencing to enhance statistical power. As before (Figure 3A), PC1 (explaining 45% of transcriptome variance) separated samples by genotype (Figure 5A). A total of 3821 genes reached the FDR adjusted p-value cut-off (<0.05) (Data 2). SHH pathway genes *GLI1, GLI2, GLI3* and *PTCH1* were significantly upregulated in *PTCH1*+/- organoids (Figure 5B), which taken together with the the effect on growth (Figure 2B) is consistent with increased SHH signal transduction. During cerebellar development, SHH signalling is most prominently known for inducing proliferation and expansion of the GCP population (Wechsler-Reya and Scott, 1999). In line with this, *PAX6* as well as *ZIC1* and *ZIC2,* two other GCP markers that are expressed throughout GC development (Aruga, 2004) and are thought to enhance SHH signalling (Brewster et al., 1998) were upregulated in heterozygous (*PTCH1+/-*) organoids. We confirmed the change in PAX6 expression was present at the protein level also using flow cytometry of day 50 organoids, which showed a significantly higher mean fluorescence intensity (MFI) in *PTCH1*+/- organoids compared to controls (p<0.01) (Figure 5C). The increased expression of specific markers of the GC population was accompanied by the increased expression of genes associated with the cell cycle and proliferation (*PCNA, MKI67, TOP2A, DLGAP5* and *CCND1*) (Figure 5B). Concurrently, early post-mitotic (*NEUROD1, NEUROD2*) and migrating GC markers (*CNTN2, CNTN1, UNC5C*) (Ackerman et al., 1997; Miyata et al., 1999; Stoykova and Gruss, 1994) were decreased. Interestingly, the increased expression of GCP markers was accompanied by the elevation of *EOMES* and *TBR1*, marking the two other main cell types derived from the RL, UBCs and glutamatergic DCN, respectively (Fink et al., 2006; Mugnaini et al., 2011). There was no evidence for differential expression of genes (*CALB2, GRM1*) associated with more differentiated UBCs (Mugnaini et al., 2011). These results may therefore suggest a broader effect of SHH-induced proliferation on RL derivatives. Together, our findings are consistent with an increased proportion of GCP, concomitant with reduced differentiation, as a result of SHH signalling-induced proliferation in *PTCH1*+/- heterozygous cerebellar organoids.

**Figure 5.**
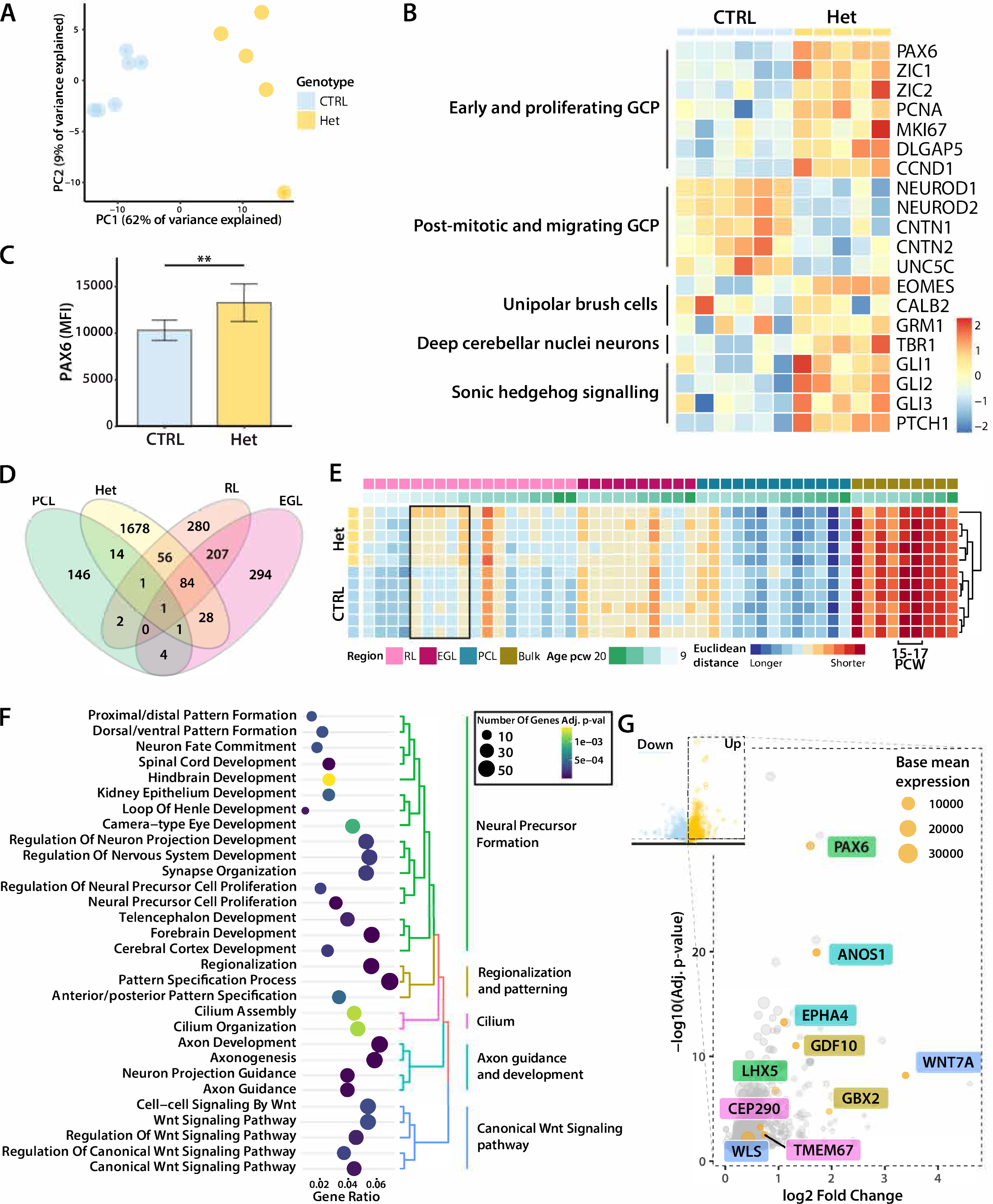
*PTCH1* heterozygous cerebellar organoids display tissue specific effects of increased SHH signalling (A) Principal component analysis using the 1000 most variable genes showing the separation of samples in principal component (PC) 1 and PC2. Control (CTRL) samples were generated with the AH017 line, Het (*PTCH1+/-*) samples with clones A3, B2, B4, and C6. (B) Expression of genes associated with different stages of rhombic lip (RL) development and RL derivatives. Only differentially expressed genes are displayed. Values are scaled within rows. The heatmap shows control (CTRL) samples generated with the AH017 line, and Het (*PTCH1+/-*) samples generated with clones A3, B2, B4, and C6. GCP: granule cell progenitor. (C) Bar graph of mean fluorescence intensity (MFI) values, measured by flow cytometry of PAX6 stained control (CTRL) and *PTCH1*+/- (Het) organoids. A total of 50,000 cells per pool were measured. n=4, **p<0.01, computed by student’s t-test. Error bars indicate standard deviation. (D) Venn diagram displaying the overlap in differentially upregulated genes. PCL: Purkinje cell layer, RL: rhombic lip, EGL: external granule cell layer. (E) Heatmap showing the Euclidean distance measurements of organoid samples with laser capture microdissected (LCM) regional samples (Aldinger et al., 2021). Samples are organised by region and age. Measurements are scaled in rows. A shorter distance relates to a higher similarity in gene expression profile. A black outline indicates the difference in similarity to RL samples seen between heterozygous and wildtype organoids. In bulk samples, the time points with the highest similarity are marked. PCL: Purkinje cell layer, RL: rhombic lip, EGL: external granule cell layer, PCW: post conception weeks (F) Gene sets uniquely enriched in heterozygous organoids. The colour of the dots corresponds to the Benjamini-Hochberg adjusted p-value, and the size of the dots corresponds to the number of genes allocated to each gene set. (G) Volcano plot of selected representative genes associated with the top uniquely differentially expressed pathways in heterozygous organoids. The size of each data point corresponds to the base mean expression of the gene.

To investigate differences between control and *PTCH1*+/- cerebellar organoids further, we sought to compare gene expression between specific regions and developmental stages of the cerebellum. To this end, organoid transcriptomes were compared with an established dataset generated from laser capture microdissected samples of the developing human cerebellum (Aldinger et al., 2021). Differentially upregulated genes marking the RL, EGL, and Purkinje cell layer (PCL) as described in the original publication, were compared to genes upregulated in *PTCH1*+/- organoids. Most differentially expressed genes in the organoid samples overlapped with RL and EGL genes (Figure 5D). Similarity with the different regional samples was then assessed by measuring Euclidean distances between LCM samples and organoid samples using the subset of spatial region defining genes. Cerebellar organoids showed the most similarity with bulk samples at post conception week (PCW) 15-17 (Figure 5E). Between the LCM samples, organoids were most similar to RL samples at 15 PCW and EGL samples at 17 PCW. Analysing distances of organoid samples to each respective region showed that *PTCH1+/-* heterozygous cerebellar organoids were more closely related to RL samples compared to control organoids. This suggests RL derivatives might make up a larger proportion of *PTCH1*+/- organoids compared to controls.

In total, 1782 differentially expressed genes were unique to *PTCH1*+/- heterozygous cerebellar organoids and not significantly altered or changed in the opposing direction in homozygous *PTCH1*-/- organoids. Gene ontology analysis was performed and revealed the enrichment of 154 gene sets, which were grouped into several main categories. Gene sets and key genes uniquely differentially expressed in heterozygous *PTCH1*+/- organoids included those involved with cerebellar neuronal precursor formation (*PAX6, LHX5*), hindbrain and cerebellar regionalisation and patterning (*GDF10, GBX2*), axon development and guidance (*ANOS1, EPHA4*), the primary cilium (*CEP290, TMEM67*), and WNT signalling (*WLS, WNT7A*) (Figure 5F,G). Taken together, these changes indicate that extensive and unique sets of genes and cerebellar developmental processes are regulated by distinct levels of SHH signalling in cerebellar organoids.

### *PTCH1*+/- cerebellar organoids display relevant features of medulloblastoma biology

The increased growth rate of *PTCH1*+/- heterozygous organoids (Figure 2) and higher expression of proliferation genes (Figure 5) is consistent with findings in the *Ptch1*+/- MB mouse model, which shows thickening of the EGL with increased and persisting proliferation of GCP prior to developing MB (Oliver et al., 2005). To confirm increased proliferation on a protein level, we analysed the expression of Cyclin B1 (CCNB1), marking cells transitioning from G2 to M phase, across cerebellar organoids from three differentiations (Figure 6A). Expression of CCNB1 was normalized to Hoechst to correct for the number of nuclei in each organoid. We found that *PTCH1*+/- heterozygous organoids expressed significantly more CCNB1 compared to control (p=0.005) (Figure 6A,B).

**Figure 6.**
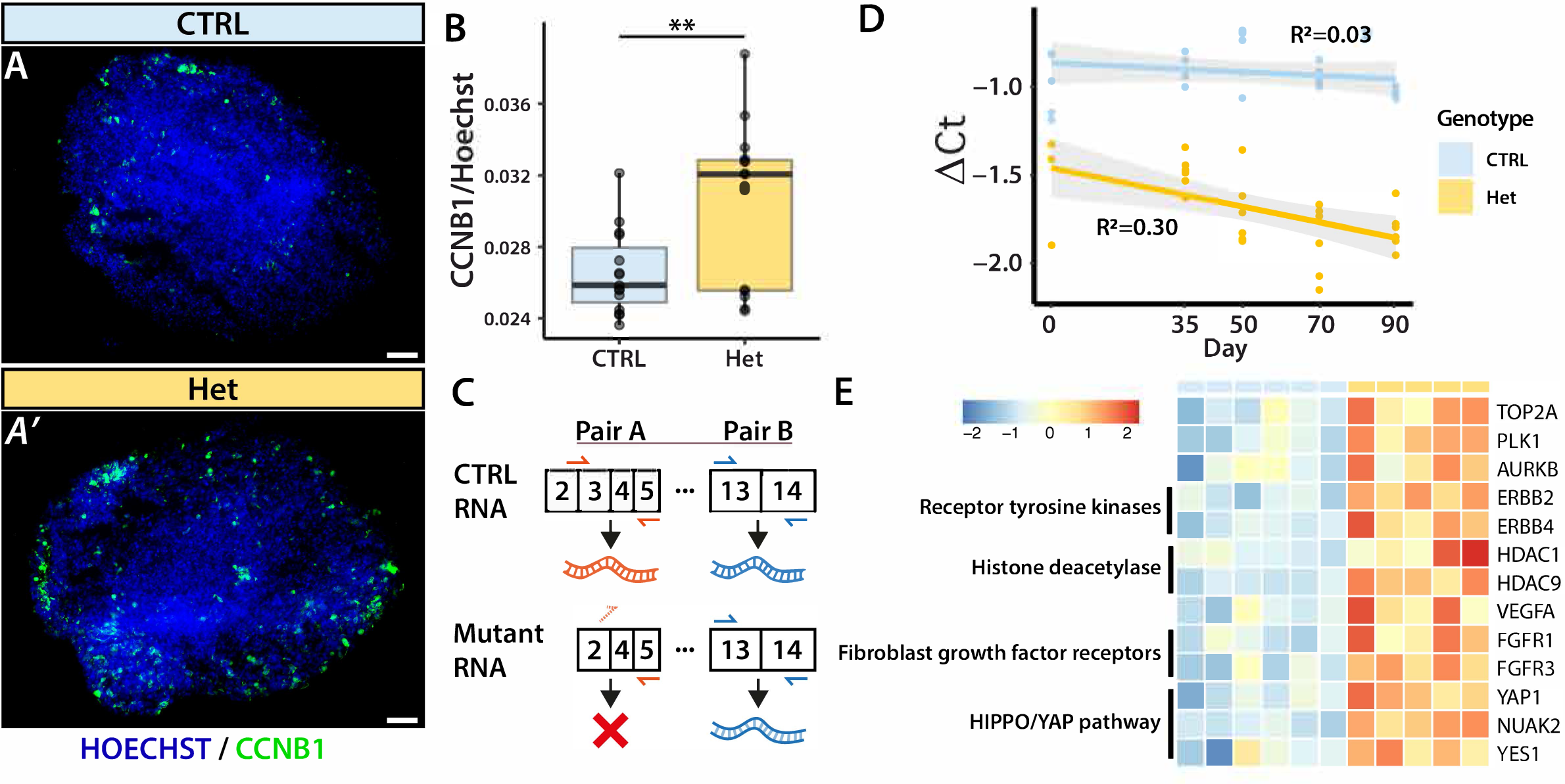
*PTCH1* heterozygous organoids display relevant features for medulloblastoma biology (A) Immunofluorescence staining of day 35 control (CTRL) and *PTCH1+/-* (Het) organoids with antibodies specific to proliferation marker CCNB1 (green). Nuclei are visualised with Hoechst. Scale bars represent 100μm. (B) Relative expression of CCNB1 normalised to Hoechst expression to correct for the number of nuclei in each organoid. Data points represent five different organoids from three differentiations. Asterisks indicate statistical significance computed by student’s t-test. **p<0.01. Error bars indicate standard deviation. (C) Schematic of priming sites of primer pair A and B. CTRL *PTCH*1 mRNA will allow amplification from both primer pairs, while mutant only allows the amplification from pair B. (D) Dot plot of ΔCt values generated by normalising the expression of pair A to pair B. Linear regression line shows the relation of the DCt with time. Lower values represent the lower relative expression of pair A, correlating with the respective expression of wildtype *PTCH1* mRNA. Data are composed of more than three biological replicates across separate differentiations. R^2^ are displayed indicating the fit of the linear regression model. (E) Example genes associated with druggable targets that were found to be upregulated in heterozygous organoids. Values are scaled within rows. The heatmap shows CTRL samples generated with the AH017 line, and Het (*PTCH1+/-*) samples generated with clones A3, B2, B4, and C6.

In addition to signs of increased proliferation, *Ptch1+/-* heterozygous mice display pre- neoplastic lesions with loss-of-heterozygosity (LOH) that frequently precede malignant transformation (Oliver et al., 2005). To examine the possible loss of *PTCH1* expression in *PTCH1*+/-heterozygous organoids, the relative expression of *PTCH1* was measured at five time points (day 0, 35, 50, 70, 90) by RT-qPCR. Mutant *PTCH1* mRNA found in *PTCH1*+/- heterozygous organoids does not contain exon 3 (Figure 6C). Thus, by comparing the amplification of a region present in both the control and gene-edited, mutant mRNA with the amplification of exon 3, the relative prevalence of *PTCH1* control mRNA can be calculated. PCR amplification from primers spanning exon-exon junction 2-3 and 5 only occurs from control *PTCH1* mRNA. Ct values generated from these primers were normalised to Ct values produced using primers that amplify exons 13 to 14 which are present in both control and mutant *PTCH1* mRNA. Linear regression showed a statistically significant age-dependent decrease of *PTCH1* mRNA in heterozygous organoids (p=0.003) but not in control organoids (Figure 6D), which might be indicative of LOH in the *PTCH1*-/- organoids.

Together, our findings show that *PTCH1*+/- heterozygous organoids display features associated with GCP proliferation and pre-neoplastic stages of MB, suggesting that the generated cerebellar organoids might be useful for the modelling of MB and as model systems to evaluate therapies. Interestingly, we found that various genes encoding known cancer targets were upregulated in *PTCH1*+/- heterozygous cerebellar organoids (Figure 6E) including MB-specific targets such as *TOP2A* (inhibited by etoposide) (Ruggiero et al., 2010; Su et al., 2022), as well as targets known in other types of cancer such as *PLK1* (volasertib), *AURKB* (barasertib)(Bavetsias and Linardopoulos, 2015), receptor tyrosine kinases such as *ERBB2* and *ERBB4* (pertuzumab, afatinib)(Hynes and MacDonald, 2009), histone deacetylases *HDAC1* and *HDAC9* (vorinostat)(Perla et al., 2020), *VEGFA* (Apte et al., 2019), fibroblast growth factor receptors *FGFR1* and *FGFR3* (Krook et al., 2021), and YAP-signalling components *YAP1*, *NUAK2*, and *YES1* (Brodowska et al., 2014; Hamanaka et al., 2019). Together, these results suggest that *PTCH1*+/- heterozygous organoids prove useful for target discovery and to guide future studies into MB driver genes in *PTCH1* mutant SHH-MB.

## Discussion

In this study, heterozygous and homozygous LOF mutations of PTCH1 were successfully introduced into iPSC from a healthy donor using CRISPR/Cas9 gene editing. The mutation caused LOF of PTCH1 but did not affect pluripotency or proliferation in iPSC. Directing these iPSC clones to cerebellar organoid differentiation demonstrated how early homozygous LOF of PTCH1 prevented cerebellar differentiation of iPSC. In contrast, organoids heterozygous for PTCH1 LOF acquired a cerebellar identity, were more proliferative, and contained an expanded glutamatergic lineage compared to control cerebellar organoids. Together, our findings support the occurrence of developmental mechanisms in cerebellar organoids that mirror *in vivo* processes of regionalisation and SHH signalling, and offer new insight into early pathophysiological events of tumorigenesis.

The pronounced effect of biallelic *PTCH1* LOF on cerebellar differentiation is in line with previous findings of mouse studies. In *Ptch1-/-* mice, neural tube closure fails and there is expansion of the ventral neural tube (Goodrich et al., 1997), which is consistent with the role of SHH signalling in orchestrating neuronal identity along the dorsoventral axis (Briscoe and Ericson, 1999; Martí and Bovolenta, 2002; Ribes and Briscoe, 2009). In vertebrates, secretion of SHH by the floor plate and notochord induces the expression of NKX2-2 in adjacent neural progenitor domains, whereas it suppresses the expression of PAX6 (Ericson et al., 1997). In line with this, *PTCH1-/-* organoids, which displayed high levels of *SHH* expression, were marked by upregulated *NKX2-2* expression and the absence of *PAX6*. Only discrete regions of the organoid expressed NKX2-2, suggesting that some level of spatial patterning occurs even *in vitro*. Furthermore inhibition of SMO using Cyclopamine rescued the expression of cerebellar markers. Therefore, in *PTCH1*-/- homozygous organoids the altered differentiation trajectory away from cerebellar identity and towards ventral fates results from hyperactivation of SHH signalling. The forebrain patterning defect is consistent with known SHH-WNT pathway interactions (Bertrand and Dahmane, 2006; Cederquist et al., 2019; Ulloa and Martí, 2010). Preventing early high level SHH signalling had enduring effects on cerebellar marker expression, which is likely to reflect progressive cellular commitment to restricted, region-specific fates (Edlund and Jessell, 1999).

Increased SHH signalling in *PTCH1*+/- organoids did not reach the threshold necessary to trigger a fate switch to ventral hindbrain identities and a cerebellar fate was retained. This enabled us to separate the effects of SHH signalling on fate specification from other SHH-regulated processes in the organoid and to gain insight into the effects of SHH signalling on the cellular constituents of the cerebellum. At an earlier stage of neural tube development when dorsal-ventral polarity is established, a high level of SHH signalling suppresses *PAX6* expression ventrally (Ericson et al., 1997). In our studies, the opposite is observed in *PTCH1*+/- heterozygous cerebellar organoids despite a higher level of SHH signalling. In the developing cerebellum, PAX6 is expressed in RL derivatives and is involved in the early migration of GCPs from the outer EGL to the inner EGL (Engelkamp et al., 1999). The EGL therefore presents a unique developmental zone where PAX6 expression persists in an environment where high levels of SHH signalling drive GCP proliferation. The co-occurring increase of *PAX6, PTCH1*, *GLI1* and *GLI2* expression in *PTCH1* heterozygous organoids suggests these organoids contain EGL-like regions that show a tissue compartment-specific effect of SHH signalling.

As GCP make up the majority of the RL lineage, our findings of increased expression of RL markers such as *PAX6* in *PTCH1*+/- heterozygous organoids may be primarily the effect of an increased GCP population. However, increased expression of *WLS*, which marks the ventricular zone of the RL that contains RL neural stem cells (Haldipur et al., 2019; Yeung et al., 2014), may also support an effect of increased SHH signalling on broader RL development. Pathologies associated with aberrant human RL development manifest by cerebellar vermis hypoplasia (CVH) (Haldipur et al., 2019). WNT signalling has been implicated as a causative factor of CVH in ciliopathies (Lancaster et al., 2011), and *PTCH1+/-* organoids also displayed increased expression of WNT pathway genes, in contrast to the *PTCH1*-/- homozygous organoids. A significant part of all CVH is caused by mutations in genes related to the primary cilium and the requirement of an intact primary cilium for SHH signalling is well described (Goetz and Anderson, 2010). The coordinated increase in gene expression related to the RL-lineage, SHH pathway, WNT pathway and cilia in heterozygous organoids is further evidence for the early involvement of SHH in human cerebellar development by regulating WNT signalling.

Our cerebellar organoid model recapitulates the genetic defect in Gorlin syndrome (also known as nevoid basal cell carcinoma syndrome) (Gorlin and Goltz, 1960), which is caused by a germline *PTCH1* heterozygous mutation and is associated with SHH-MB. Furthermore, the increased organoid growth, upregulation of cell cycle genes, and increased expression of mitotic markers found in heterozygous organoids resemble the preneoplastic stage in *Ptch1+/-* mice (Oliver et al., 2005). *Ptch1+/-* mice exhibit a persisting hyperplastic EGL with pre- neoplastic lesions that have lost the wildtype allele. Similarly, wildtype *PTCH1* mRNA levels decreased in heterozygous organoids, suggesting loss-of-heterozygosity (LOH), which is frequently seen in both mouse models of MB (Ishida et al., 2010; Pazzaglia et al., 2006), and human *PTCH1*-mutant MB (Northcott et al., 2017; Tamayo-Orrego et al., 2020). Studies have shown a clear association of SHH activation-induced DNA replication stress and elevated homologous recombination (Tamayo-Orrego et al., 2020). In line with these findings, heterozygous organoids displayed signs of loss of wildtype *PTCH1,* suggesting our *in vitro* model could be used to gain further insights into these events.

We have shown that cerebellar organoid differentiation can accurately capture relevant human developmental phenotypes *in vitro,* and complement findings from mouse models. This is especially relevant for MB where findings in mouse models (Tamayo-Orrego et al., 2016) can diverge from large human genomic studies (Kool et al., 2008; Northcott et al., 2011; Skowron et al., 2021). Future investigations can apply the latest findings from these genome studies to cerebellar organoids to develop novel, human-specific models that are poised to aid the discovery of much needed new treatments for MB patients.

## Experimental Procedures

### Resource availability

Further information and requests should be directed to corresponding author Esther Becker.

Materials availability: There are restrictions to the availability of iPSC lines due to the lack of an external centralized repository for its distribution and our need to maintain the stock. We are glad to share cells with reasonable compensation by requestor for its processing and shipping. We may require a payment and/or a completed Materials Transfer Agreement if there is potential for commercial application.

Data and code availability: All data and code is available on https://github.com/mxvssn

### Induced pluripotent stem cell line and culture

The previously described human iPSC line AH017-3 was used (Handel et al., 2016). iPSC were cultured on hESC-qualified Matrigel matrix (Corning®, Cat#354277) in mTeSR1 medium (Stemcell Technologies, Cat#85850) or OXE8 medium (2mM GlutaMax, 0.1*μ*g/mL Heparin, 0.22mM Ascorbic Acid, 15mM HEPES pH7.4, 100ng/mL FGF2, 2ng/mL TGF*β* in Advanced DMEM/F12) (Vaughan-Jackson et al., 2021). The culture medium was changed daily, and passaged using 0.5mM EDTA in PBS. 10*μ*M Rho kinase inhibitor (Ri) Y-27632 (Abcam, Cat#ab120129) was added to the culture medium during the first 24 hours after passaging. Cells were kept in culture no longer than five weeks to minimise the chance of karyotypic changes. For each new experiment, a vial was used from the original master stock. This stock had passed quality control including single nucleotide polymorphism (SNP) genotyping and mycoplasma testing.

### CRISPR-mediated Gene Editing

Guide RNAs (gRNAs) were designed using CCtop (Stemmer et al., 2015). The exonic gRNA GT GTT GTA GGA GCG CTT CTG and intronic gRNA GA TTT ATC GTT TCT CGA GTT were picked based on their predicted specificity and efficiency. Predicted off-target sites are listed in Figure S4.

crRNAs and tracrRNAs were heated to 95°C before slowly cooling to RT to generate crRNA/tracrRNA hybrids. The exonic and intronic targeting crRNA/tracrRNA hybrids were then mixed together in a 1:1 ratio. Ribonuclease proteins were made by complexing 44μM crRNA/tracrRNA hybrids with 36μM HiFi Cas9 Nuclease V3 (IDT, Cat#1081060). 1μL of ribonuclease proteins was mixed with 9μL iPSC suspension (200,000 cells total) and the mixture was electroporated using the NeonTM Transfection System (Cat#MPK10096; HiTrans 1400V 20ms width, one pulse). The cell pool was cultured to produce a CRISPR-pool stock and allow genomic DNA extraction (DNeasy kit, Qiagen, Cat#69504). Successful CRISPR editing was visualized by PCR amplification of the target region (primers listed in Table S1). The mutant product was expected to be 243bp shorter compared to the wildtype sequence. Monoclonal iPSC stocks were generated by single-cell plating on mouse embryonic feeders. Clones derived from a single cell were microscopically picked using a wide bore P200 tip. Selected clones were expanded to master stocks that were used for all downstream experiments.

### Cerebellar organoid differentiation

Cerebellar organoids were generated as described (van Essen et al., 2022; Nayler et al., 2021; Watson et al., 2018). In brief, iPSC were detached using TrypLE™ Express Enzyme (1X) (Gibco™, Cat# 12604013) and resuspended in Induction Medium (50% Iscove’s Modified Dulbecco’s Medium (Gibco™, Cat# 31980022), 50% Ham’s F-12 Nutrient Mix (Gibco™, Cat# 31765027), 7*μ*g/mL insulin (Sigma, Cat# I1882), 5mg/mL BSA (|Sigma, Cat# A3156-5G), 1% Chemically Defined Lipid Concentrate (Gibco™, Cat# 11905031), 450*μ*M 1-thioglycerol (Sigma, Cat# M-6145), 15*μ*g/mL apo-transferrin (Sigma, Cat# T1147), 1% Penicillin/Streptomycin (Gibco™, Cat#15140122)) containing 50μM Rho-kinase inhibitor(Abcam, Cat#ab120129) and 10μM SB431542 (Tocris, Cat#1614/10). Embryoid bodies (EBs) were made by seeding cells to 10,000 cells per well in a low-attachment V-bottom 96-well plate (Greiner Bio-one, Cat# 651970) and incubating at 37°C with 5%CO2. Two days after seeding, cerebellar lineage induction was started using by supplementing the medium with FGF2 (R&D Systems, Cat#4114-TC-01M) to final concentration of 50ng/mL. Subsequent medium changes were performed weekly. On day 7, one-third of the medium was changed, and on other days the medium was changed in full. On day 14, organoids were transferred to low-attachment 48-well plates (Greiner Bio-one, Cat#677970). On day 21, the culture medium was changed to Differentiation medium (Neurobasal® medium (Gibco™, Cat# 21103049), 1% GlutaMAXTM (Gibco™, Cat# 35050061), 1% N2 supplement (Gibco™, Cat# 17502048), 1% Penicillin/Streptomycin (Gibco™, Cat# 15140122)). Long-term culture from day 35 onward was performed on PTFE 0.4-μm pore size transwell membranes (Millicel, Cat#PICM0RG50).

For cyclopamine treatment, embryoid bodies of control and *PTCH1-/-* iPSC were made as above. After two days, treatment was started and either DMSO, 1*μ*M Cyclopamine, 5*μ*M Cyclopamine, or 500nM SAG was added to the medium upon medium change. Organoids were harvested after 35 days for RNA extraction and RT-qPCR or immunostaining.

### RT-qPCR

RNA was isolated using the RNeasy Plus mini kit (Qiagen, cat# 74034) following the manufacturer’s protocol. Three replicates extracted from different passages were collected per clone. Data shown in the figures was generated with the AH017 control line (CTRL), the *PTCH1+/-* A3 line (Het) and/or the *PTCH1-/-* H3 line (Hom). RNA was reverse transcribed into cDNA using the SuperScript™ III First-Strand Synthesis System (Invitrogen™, Cat#18080051). Standard curves to determine primer efficiency and optimal cDNA input were run prior to the experiment. RT-qPCR was performed using Fast SYBR Green Master Mix (Applied Biosystems, Cat#4385612) on the Applied Biosystems StepOne Plus qPCR machine with primers designed using Primer3Plus. Primers used are listed in Table S1. Expression of genes of interest (‘GOI’) was normalised to the expression of *ACTB* and *GAPDH* (‘REF’) to generate delta-CT values (*ΔCT* = *CT*^*GOI*^ − *CT*^*REF*^). The average of these two delta-CT values was used for statistical analysis and visualisation. In comparisons with a control group, *ΔΔ*CT values were calculated (*ΔΔCT* = *ΔCT*^*Test*^ − *ΔCT*^*Control Averagr*^). The expression of reference genes in each experiment was subjected to statistical testing for changes associated with the different conditions.

### Immunostaining

Organoids were fixed in 4% paraformaldehyde (PFA) and embedded in Optimal Cutting Temperature compound to allow cryosectioning. Cryosections were permeabilised in 0.3% Triton-X100 (Sigma, Cat#X100-100ML) in PBS (PBST) and blocked with 2% skim milk (Thermofisher, Cat#LP0031B) in PBST (blocking buffer). Primary antibodies (Table S2) diluted in blocking buffer were incubated overnight. Secondary antibodies (Table S3) diluted in blocking buffer were applied for two hours. Nuclei were visualized with Hoechst (Thermofisher, Cat#62249) and sections were mounted in anti-fading mounting solution (Vectashield, Cat#H-1000-10). Immunostained cryosections were imaged using a Zeiss Axioplan 2 widefield microscope or an Olympus FV1000 laser scanning confocal microscope. Data shown in the figures was generated with the AH017 control line (CTRL), the *PTCH1+/-* A3 line (Het) and/or the *PTCH1-/-* H3 line (Hom).

### Flow Cytometry

Organoids were washed once in PBS at room temperature and enzymatically digested using Neuron Isolation Enzyme (ThermoFisher, Cat#88285). Cells were washed in FACS buffer (1% BSA in PBS) and fixed in 4% PFA. Permeabilisation was achieved using FACS buffer with 0.1% saponin. Cells were incubated with PAX6 antibodies conjugated to APC (lightning link® conjugation kit, Abcam, Cat#ab201807) for 30 minutes. Cells were washed once more in FACS buffer and stained with Hoechst. Flow cytometry was performed on the FACS Canto BD (BD Biosciences). Mean fluorescent intensity of 50,000 cells was measured and statistical analysis was performed as described above.

### RNA Sequencing

RNA was isolated using the RNeasy Plus Micro kit (Qiagen, Cat# 74034). RNA integrity was measured using the RNA 6000 Pico kit (Agilent, Cat# 5067-1513) on the 2100 Bioanalyzer instrument (Agilent). Samples that passed this quality measurement (RIN > 9) were submitted for RNA sequencing. RNA concentration was determined using Qubit™ 4 Fluorometer following the manufacturer’s protocol. A total of 400ng per sample was submitted for library preparation. Library preparation and sequencing were performed by Novogene Co Ltd. Each library was submitted to 25 million paired-end reads (50M total). Gene transcripts were quantified using Salmon 15.2 with GC content bias and sequencing length bias correction. Transcript quantifications were imported in ’*R* programming software’ using ’tximeta’ (version 1.14.1) (Ensembl Homo sapiens release 97). Differential gene expression was performed using ’DESeq2’ (version 1.36.0) (Love et al., 2014) and the *lfcshrink* function that uses Bayesian shrinkage estimators for effect sizes (Zhu et al., 2019). Differentially expressed genes were identified as having a Benjamini-Hochberg adjusted p-value below 0.05. Gene expression was normalised using Variance-stabilising transformation (VST) (’vsn’ package version 3.64.0). Principal component analysis (PCA) was performed using prcomp (’stats’ package version 4.2.1) using the top 1000 most variable genes and visualised using ’ggplot2’ (version 3.3.6). Gene expression was visualised using ’pheatmap’ (version 1.0.12). Gene set variation analysis (GSVA) was performed using the GSVA package (version 1.44.2). Gene ontology analysis was performed using the Clusterprofiler package (version 4.4.4).

### Statistics

Statistical analysis was performed in *R* programming software. Statistical difference between more than two groups was determined using ANOVA with Dunnett’s post-hoc test or Tukey’s post-hoc test. Statistical differences between two groups were performed by student’s t-test.

P-values were corrected for multiple testing using Benjamini-Hochberg Procedure. An adjusted p-value below 0.05 was considered significant. Error bars represent standard deviation (SD) unless otherwise indicated.

## Acknowledgments

We would like to thank P. Kilfeather for assisting with the alignment of bulk RNA sequencing data. This work was supported by Cancer Research UK (CRUK) grant number C2195/A28699, through a CRUK Oxford Centre Clinical Research Training Fellowship.

## Author contributions

M.E, J.R., S.C., J.J.and E.B. conceptualized the study and designed the experiments. M.E. performed the experiments and data analysis. M.E. drafted the paper and made the figures. M.E., S.C., J.J. and E.B. co-authored and edited the manuscript. All of the authors read, edited, and approved the final version of the manuscript.

## Datasets

Data 1: Table of differentially expressed genes in homozygous clones

Data 2: Table of differentially expressed genes in heterozygous clones

