## Supplemental Information for "*PTCH1* mutant human cerebellar organoids are associated with altered neural development and early pathways of medulloblastoma oncogenesis"

**Supplemental Items**

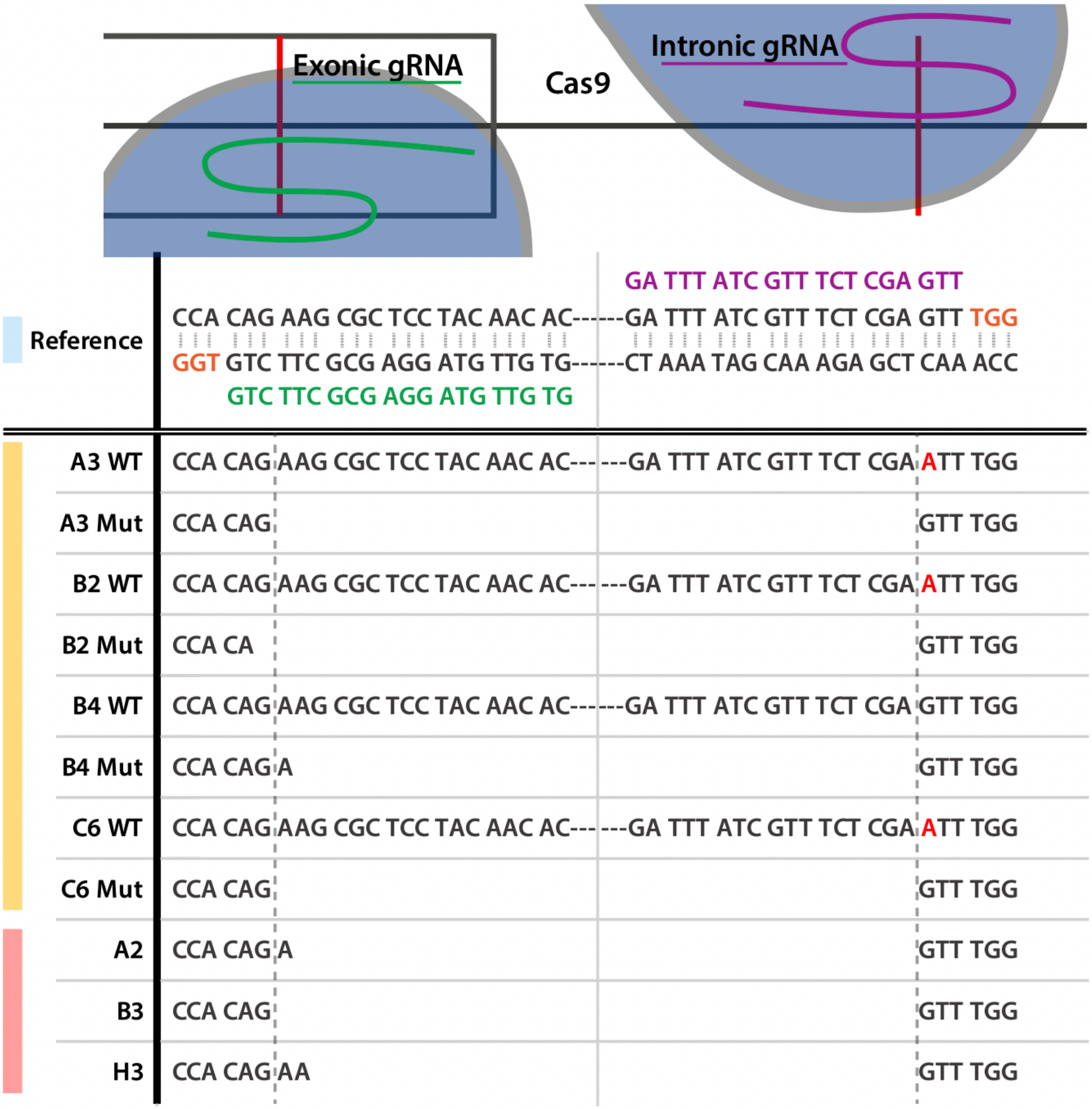

**Figure S1 Sanger sequencing of CRISPR target site in the *PTCH1* gene (related to Figure 1)**
Top: Sequencing of control *PTCH1* sequence (reference). Guide RNA (gRNA) sequences (exonic in green, intronic in purple) and respective protospacer adjacent motifs (marked orange in reference sequence) are displayed with the coding strand of the reference sequence on top. Below: Sequencing of mutant clones. The expected cut sites are depicted as vertical dashed lines. Sanger sequencing revealed uniform cutting at the intronic target site compared to more heterogeneous cutting at the exonic target site. The genomic DNA region between the target sites is absent in the mutant amplicons (mut). Point mutations at the intronic target site resulting from the Cas9-induced double strand break are detected in heterozygous clones (G>A) (marked in red).

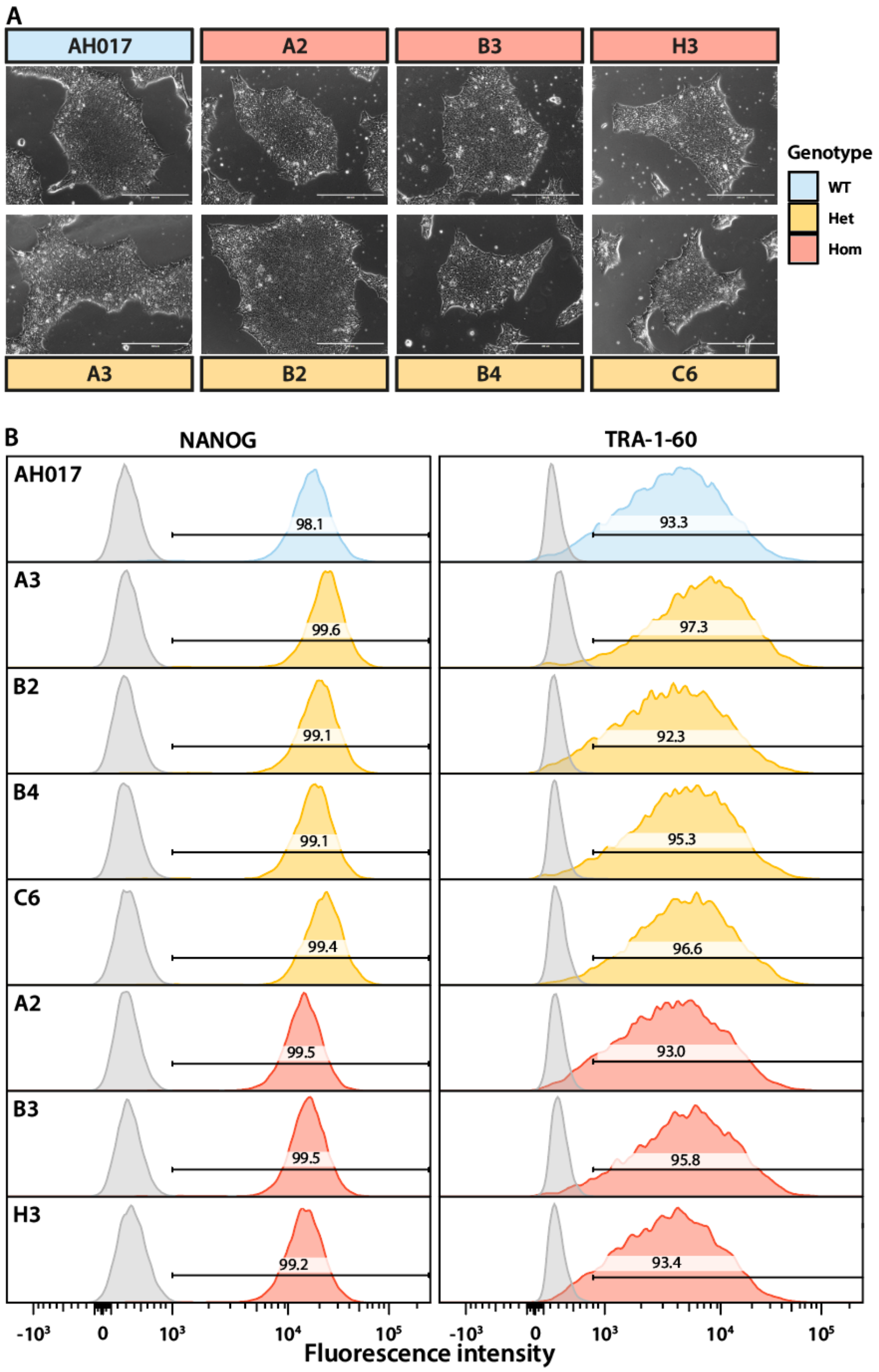

**Figure S2 PTCH1 mutant clones maintain iPSC morphology and pluripotency marker expression (related to Figure 1)**

A) Brightfield images of iPSC clones. Bars are 1000µm. All clones display normal iPSC morphology with large nucleus-to-cytoplasm ratio and growth in densely packed colonies.

B) A total of 50,000 cells were measured by flow cytometry. Density plot shows the distribution of fluorescence intensity along the x-axis. Coloured histograms represent cells stained with antibodies targeting NANOG (left) or TRA-1-60 (right). Grey histograms represent cells stained with isotype control antibodies to assess non-specific binding and autofluorescence. The percentage of cells with fluorescence intensities above the isotype control condition are depicted.

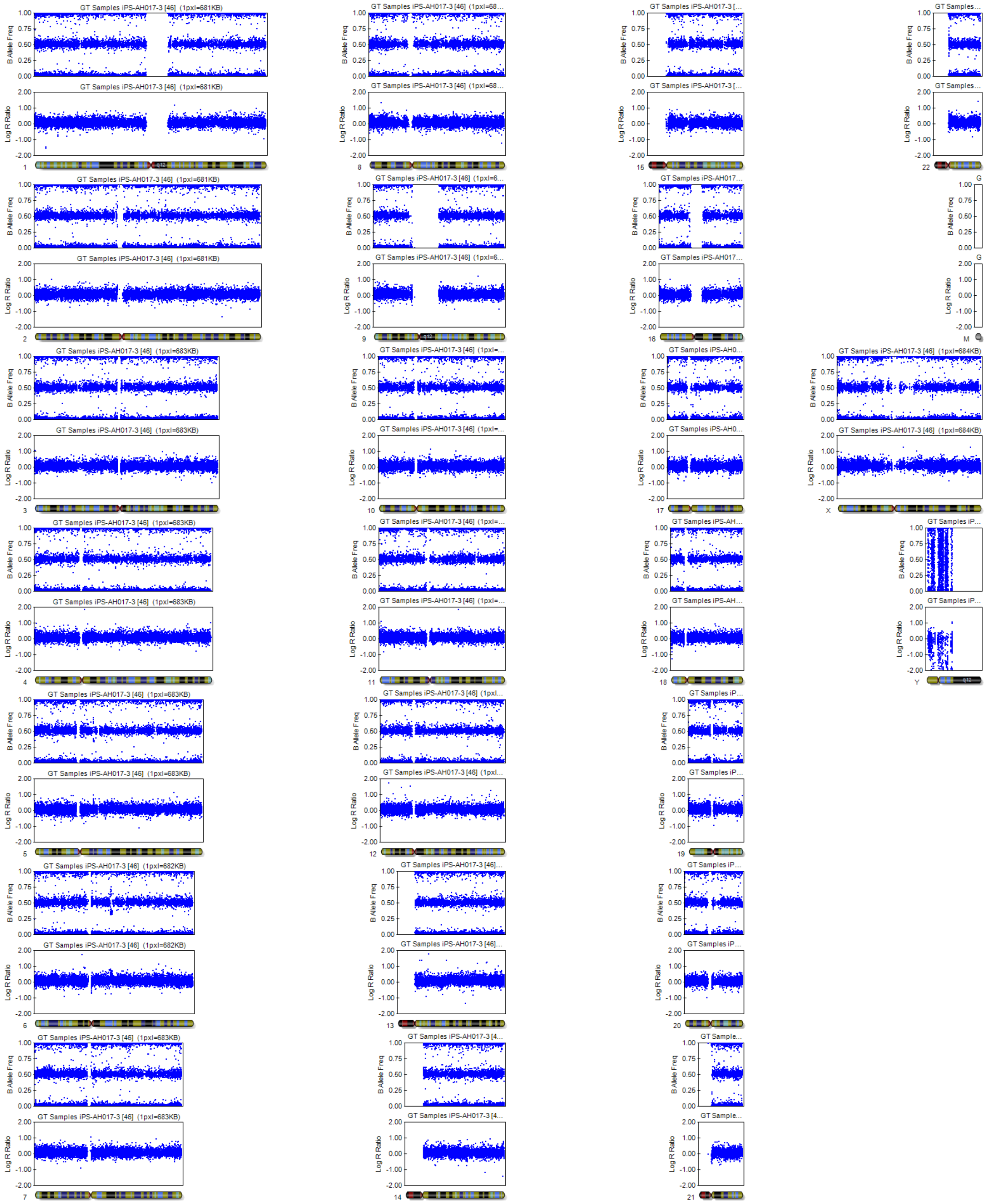
GT Samples iPS-AH017-3 [46] (1pxl=681KB)
GT Samples iPS-AH017-3 [46] (1pxl=683KB)
GT Samples iPS-AH017-3 [46] (1pxl=682KB)
GT Samples iPS-AH017-3 [46] (1pxl=684KB)
B Allele Freq
Log R Ratio
X
Y
M

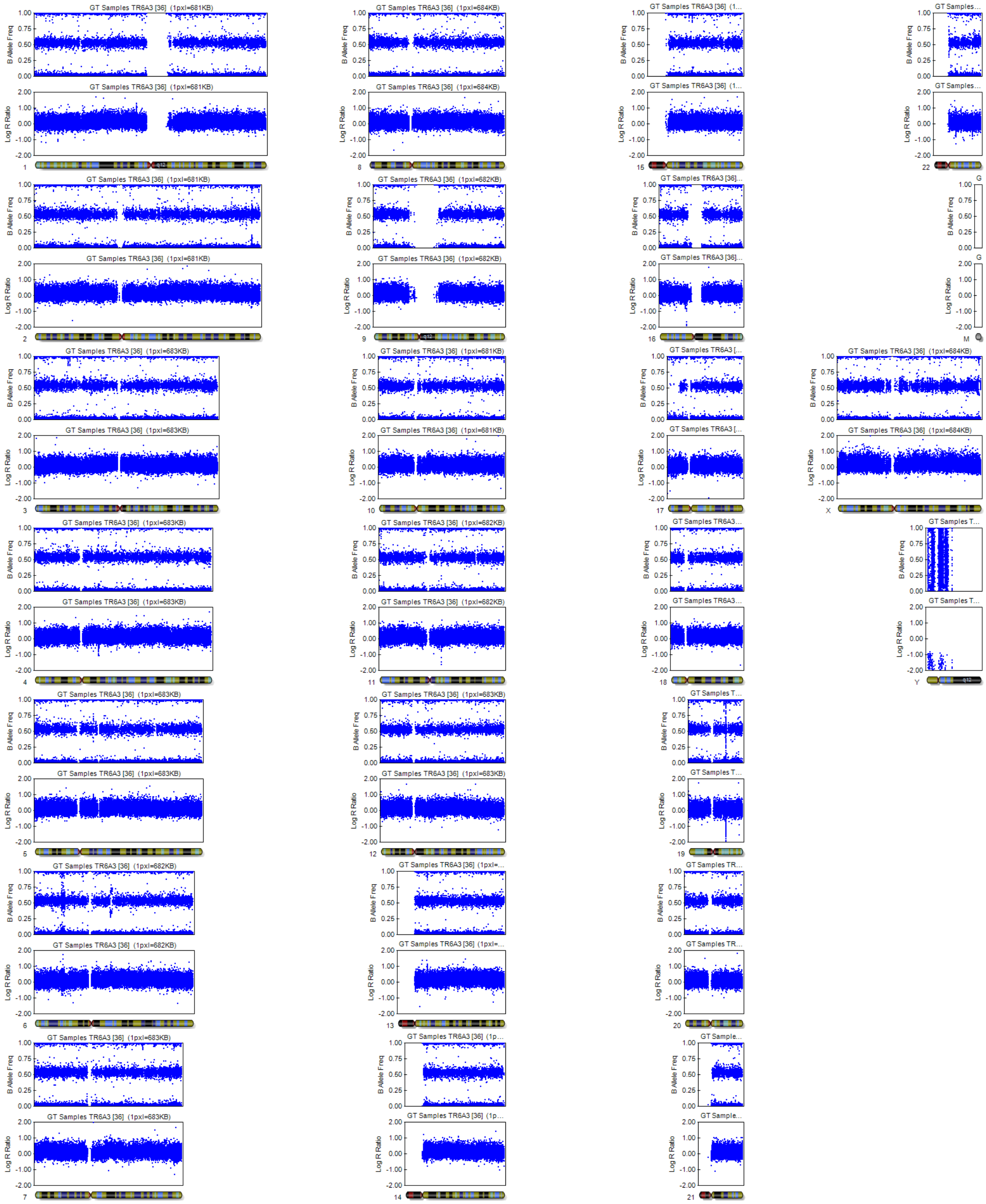
GT Samples TR6A3 [36] (1pxl=681KB)
GT Samples TR6A3 [36] (1pxl=684KB)
GT Samples TR6A3 [36] (1pxl=682KB)
GT Samples TR6A3 [36] (1pxl=683KB)
B Allele Freq
Log R Ratio
1.00
0.75
0.50
0.25
0.00
2.00
1.00
0.00
-1.00
-2.00
X
Y
M

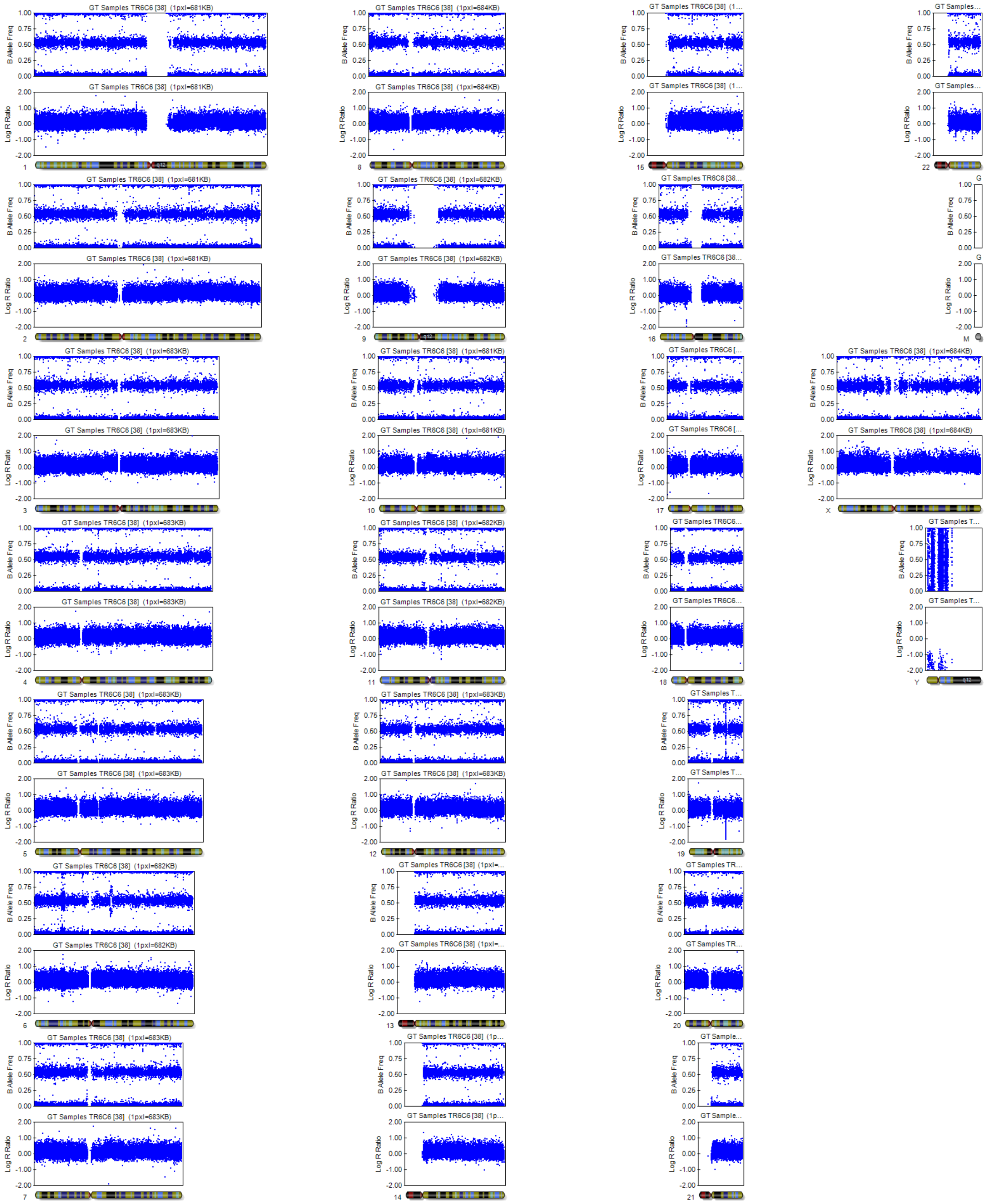
GT Samples TR6C6 [38] (1pxl=681KB)
B Allele Freq
Log R Ratio
X
Y
M

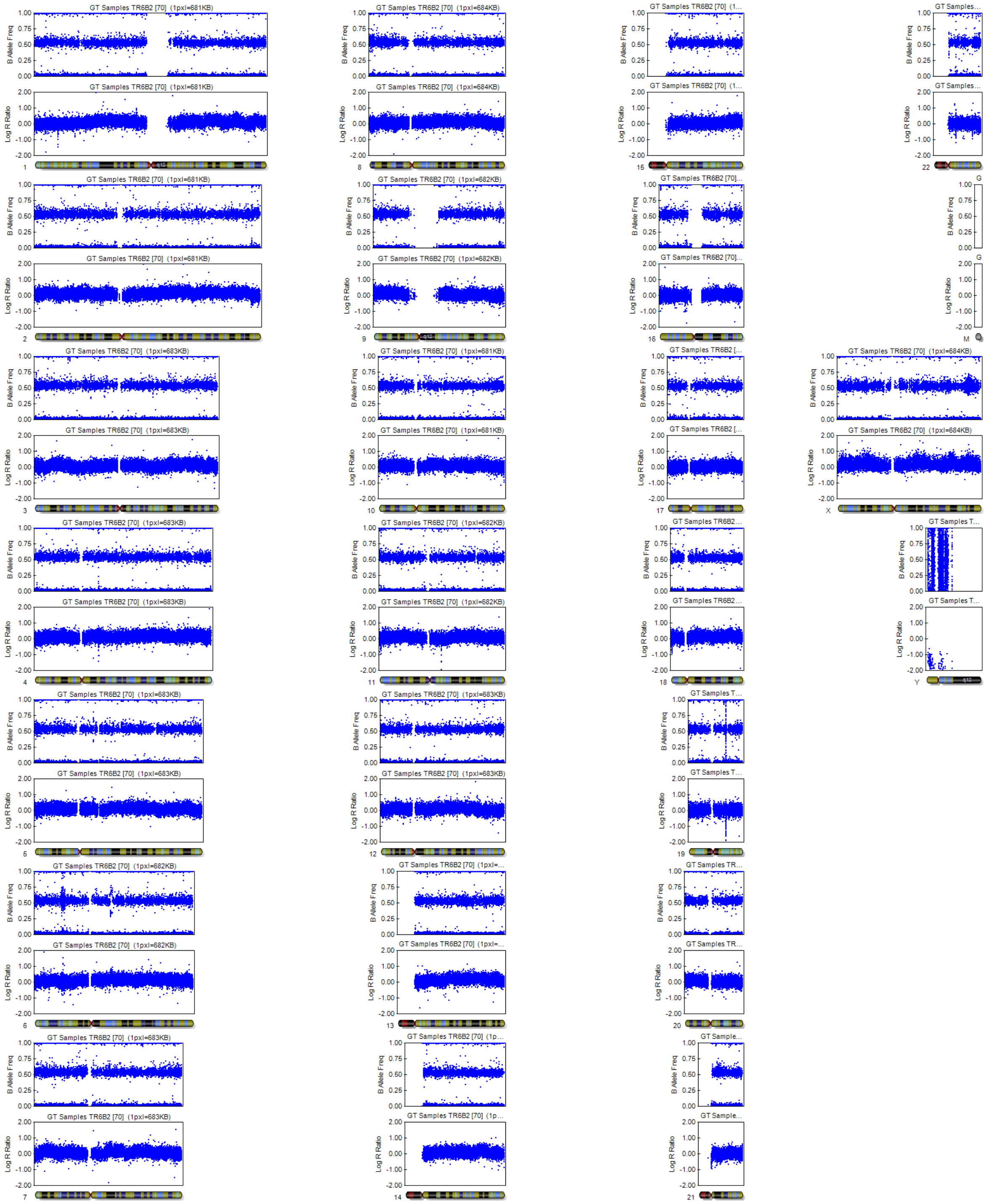

GT Samples TR6B2 [70] (1pxl=681KB)
GT Samples TR6B2 [70] (1pxl=682KB)
GT Samples TR6B2 [70] (1pxl=683KB)
GT Samples TR6B2 [70] (1pxl=684KB)
B Allele Freq
Log R Ratio
1.00
0.75
0.50
0.25
0.00
2.00
1.00
0.00
-1.00
-2.00
X
Y
M
q12

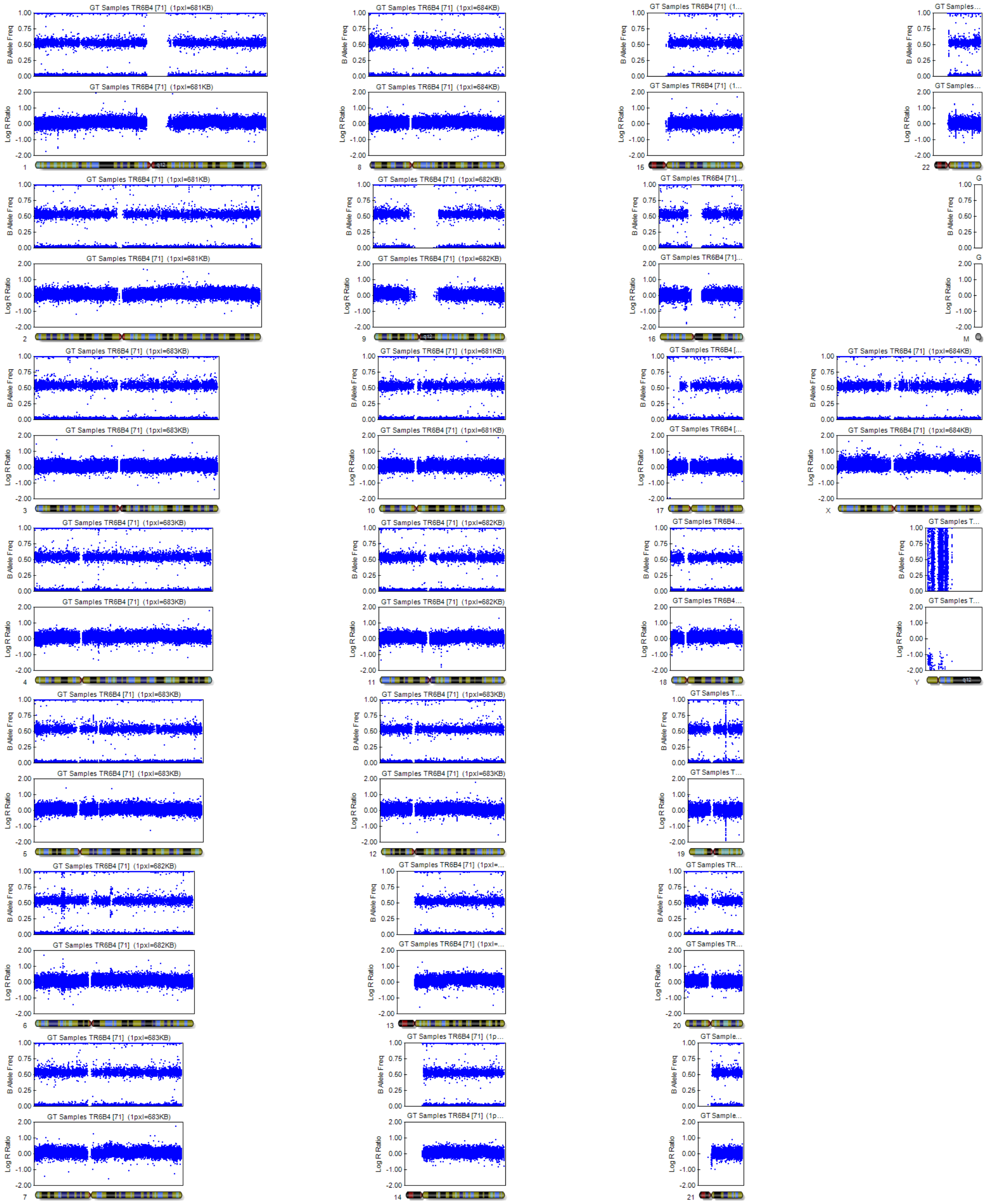

GT Samples TR6B4 [71] (1pxl=681KB)
GT Samples TR6B4 [71] (1pxl=684KB)
GT Samples TR6B4 [71] (1pxl=682KB)
GT Samples TR6B4 [71] (1pxl=683KB)
B Allele Freq
Log R Ratio
X
Y
M

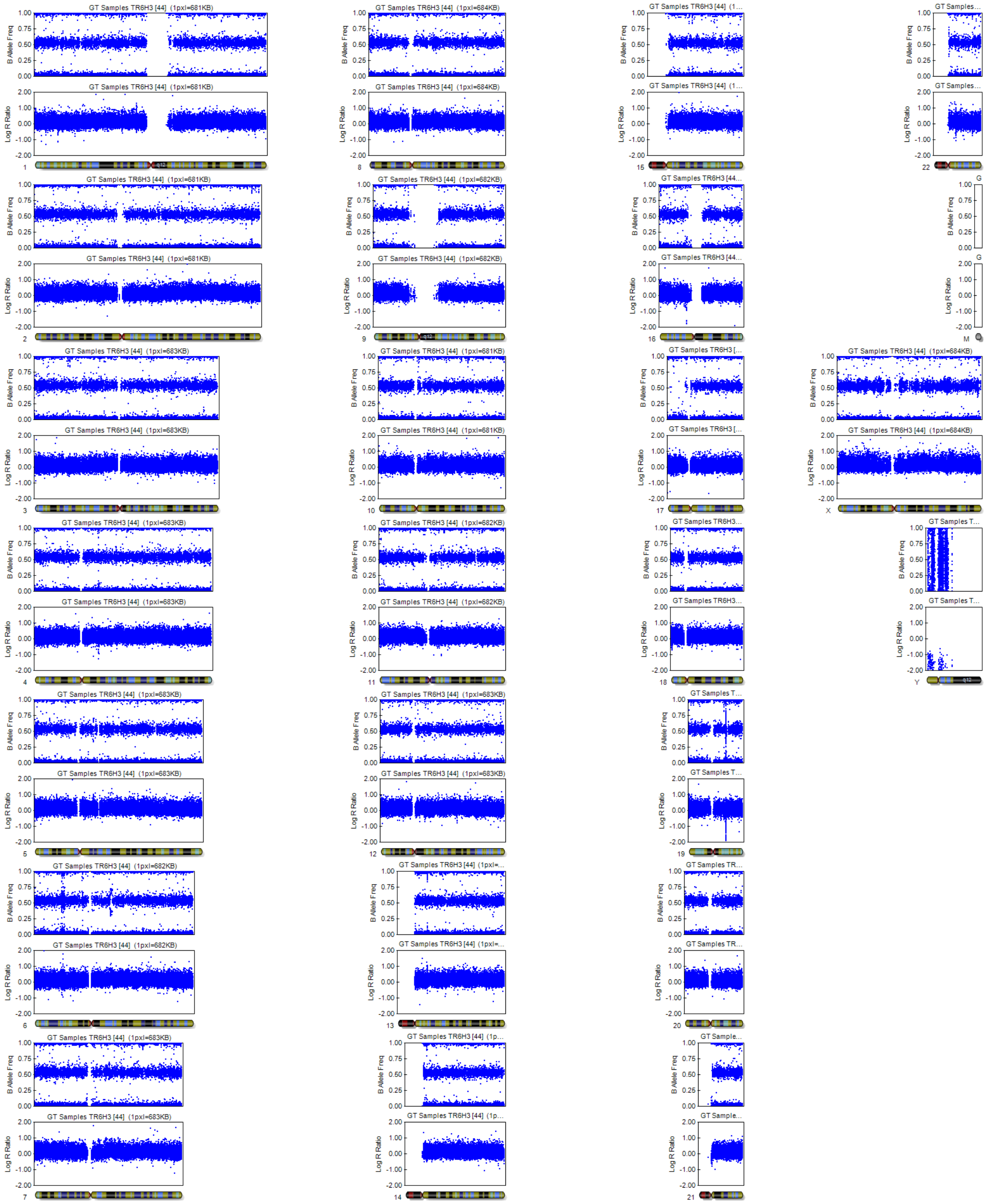
GT Samples TR6H3 [44] (1pxl=681KB)
GT Samples TR6H3 [44] (1pxl=684KB)
GT Samples TR6H3 [44] (1pxl=683KB)
GT Samples TR6H3 [44] (1pxl=682KB)
B Allele Freq
Log R Ratio
X
Y
M

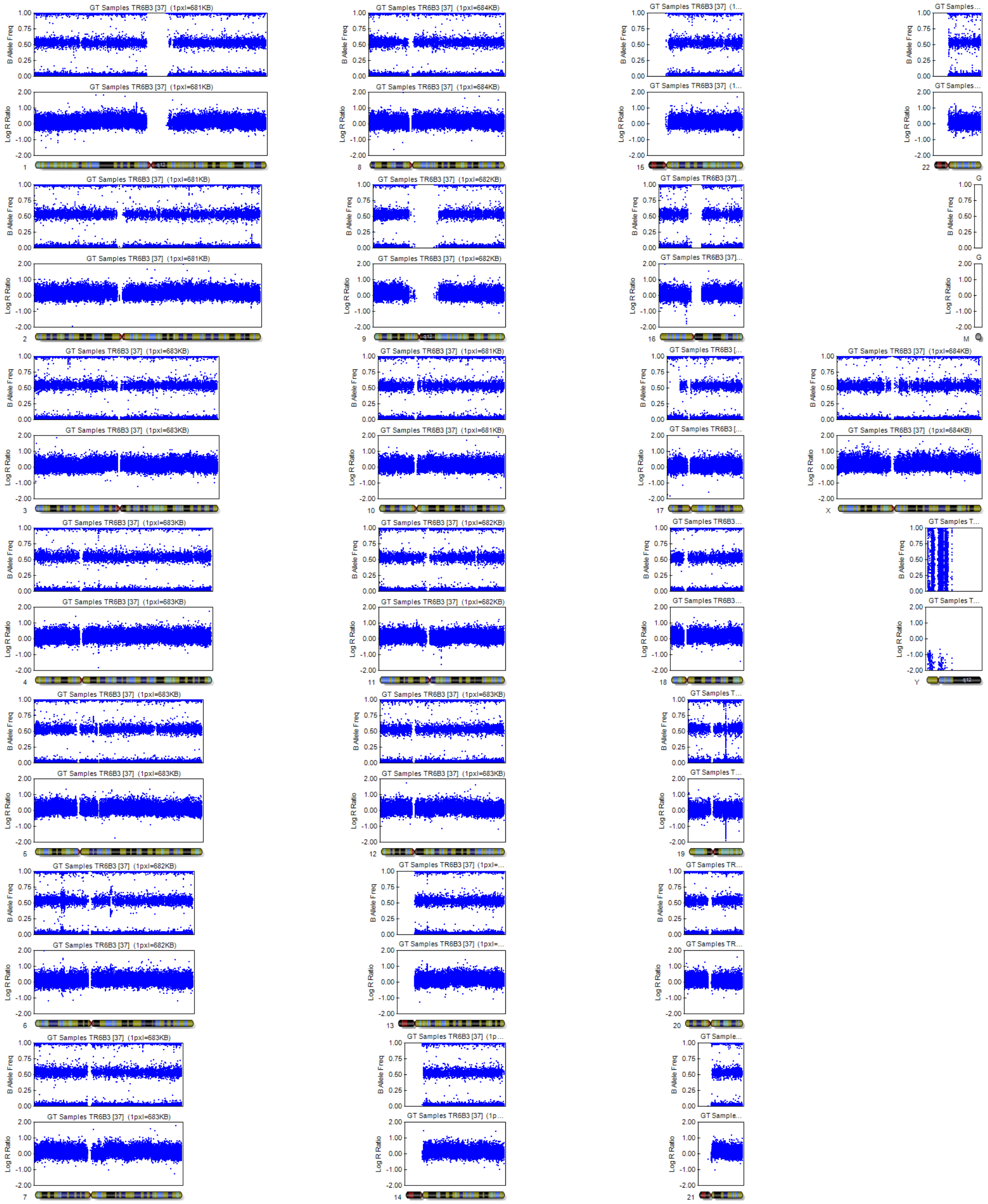
GT Samples TR6B3 [37] (1pxl=681KB)
GT Samples TR6B3 [37] (1pxl=684KB)
GT Samples TR6B3 [37] (1pxl=682KB)
GT Samples TR6B3 [37] (1pxl=683KB)
B Allele Freq
Log R Ratio
X
Y
M

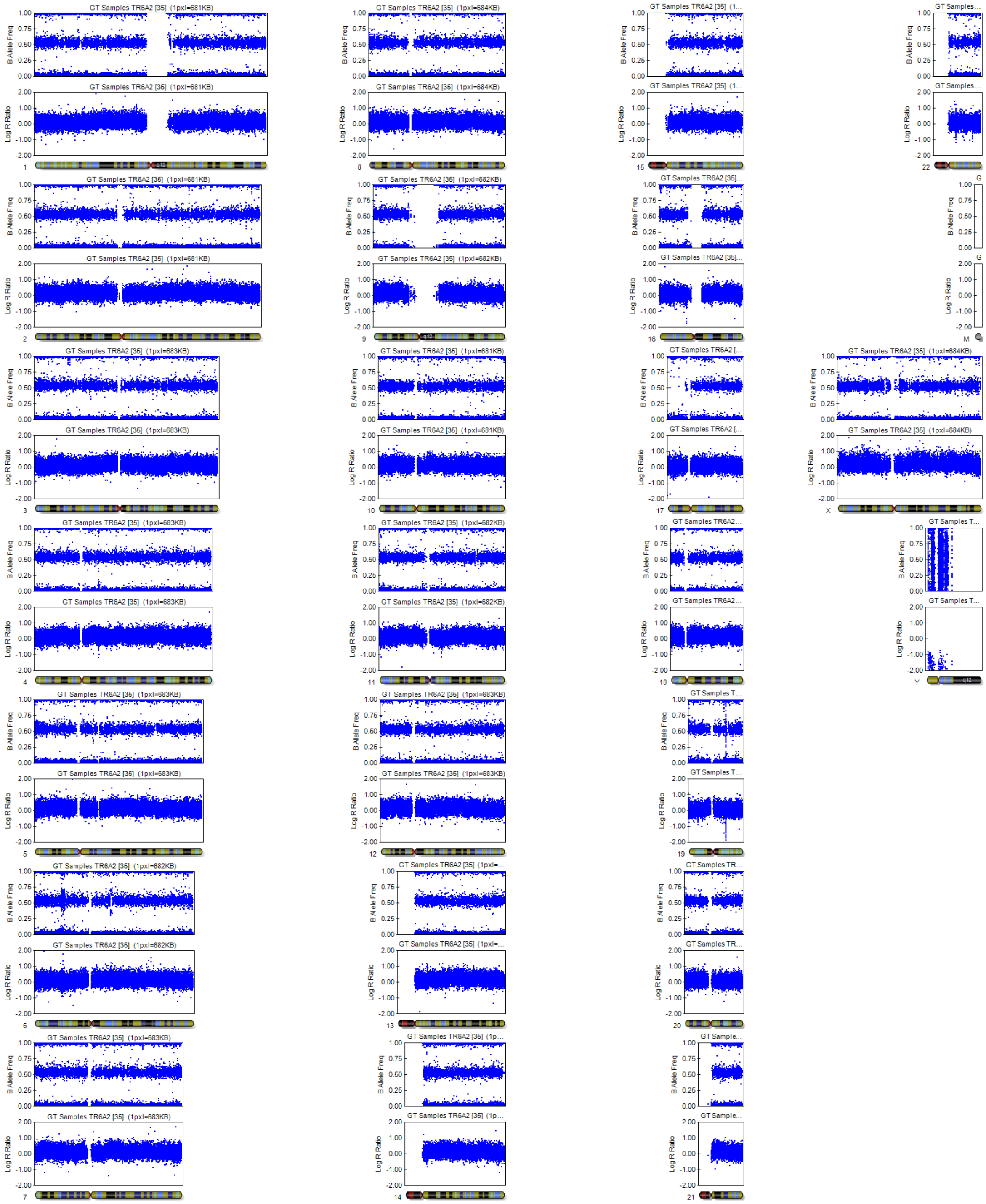

**Figure S3 SNP array to test for chromosomal aberrations in gene edited iPSC clones (related to Figure 1)**
For each chromosome, the first dot plot displays the B-allele frequency and indicates whether the SNV is heterozygous (data points fall at around 0.5) or homozygous (data point at around 0.0 of 1.0). The second plot displays the log R value (first dot plot, indicated with R on the left side) is given, representing the probe intensity of individual SNVs. Chromosomal gains are identified by doubling of the log R value while halving the R value indicates loss. The different clones are given in the following order: control line: AH017-3; heterozygous (*PTCH1+/-*) clones: A3, C6, B2, B4; homozygous (*PTCH1-/-*) clones: H3, B3, A2.

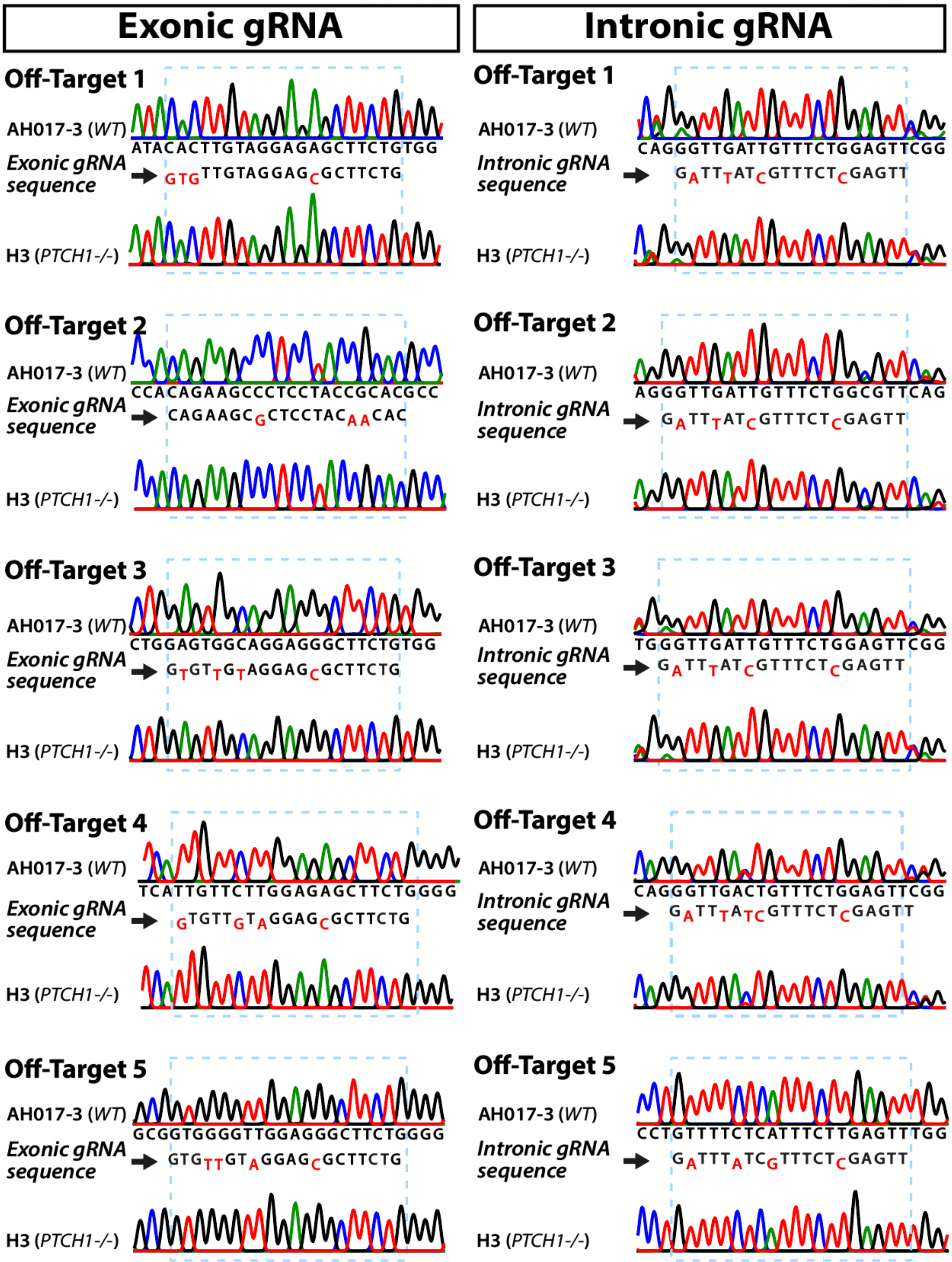

**Figure S4 Sanger sequencing does not reveal off-target CRISPR-Cas activity (related to Figure 1)**

Sanger sequencing of the top five off-target sites of each guide RNA. The alignment with each respective gRNA is shown below the wildtype strand with mismatches shown in red. Sequence trace of both the wildtype and tested homozygous mutant (H3) were identical.

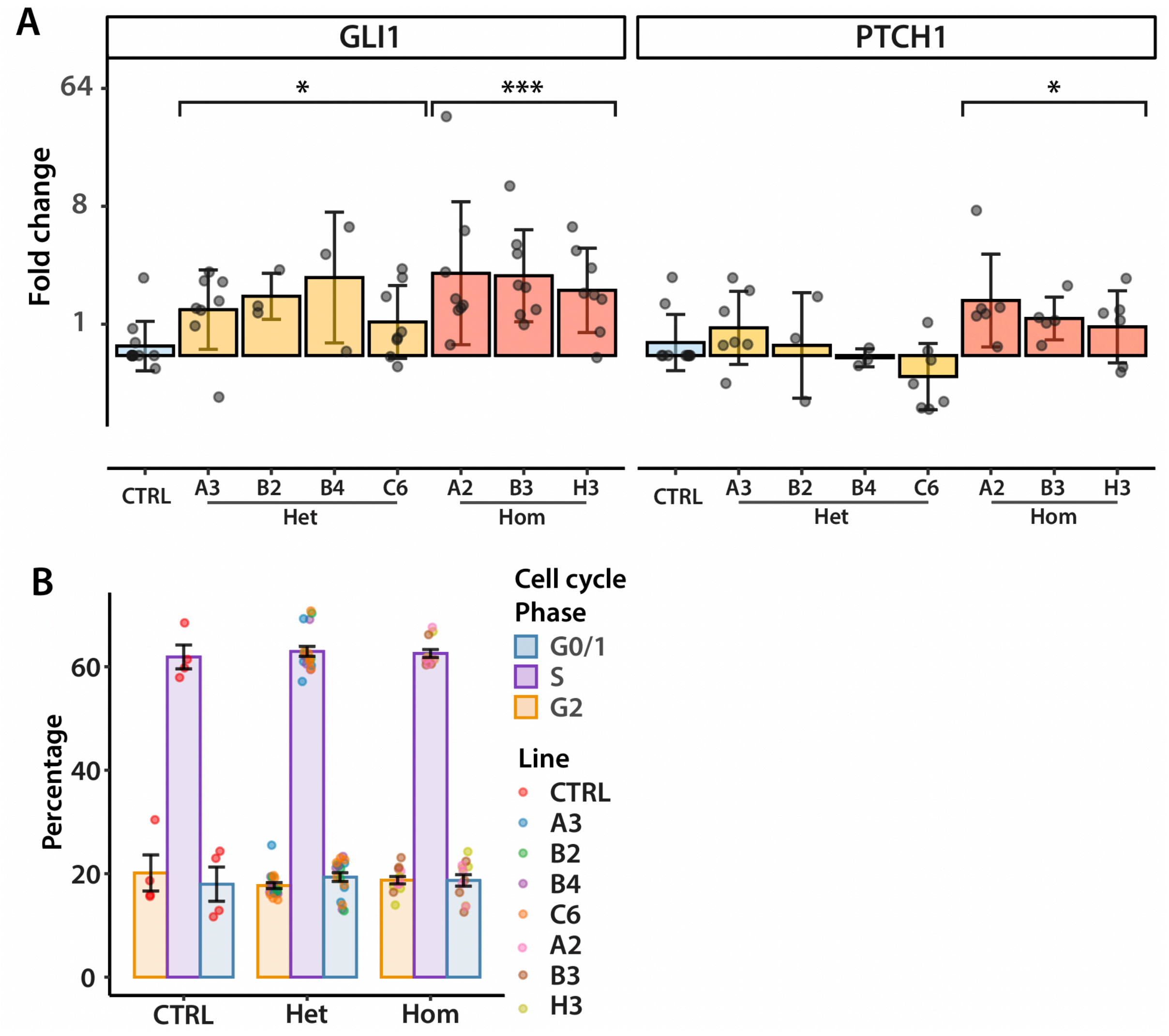

**Figure S5 Midbrain-hindbrain markers are expressed by *PTCH1* mutant organoids (related to Figure 1)**

(A) Messenger RNA expression of *GLI1* and *PTCH1* as measured by RT-qPCR. Expression is relative to the parent line (CTRL) and normalized using *ACTB* and *GAPDH*. Statistically significant differences on the genotype level are calculated on the ΔΔCt values by one way ANOVA with Dunnett's post hoc test. Significance is indicated by asterisks. *$p<0.05$, ***$p<0.001$. Error bars represent standard deviation. Data points are generated from independent passages. n>3 for all clones.

(B) Proportions of cells in G0/1 phase (orange), S phase (purple) and G2 phase (blue) as percentage of whole. Data points are generated from independent passages. n=3 for all clones. Error bars indicate standard error of the mean.

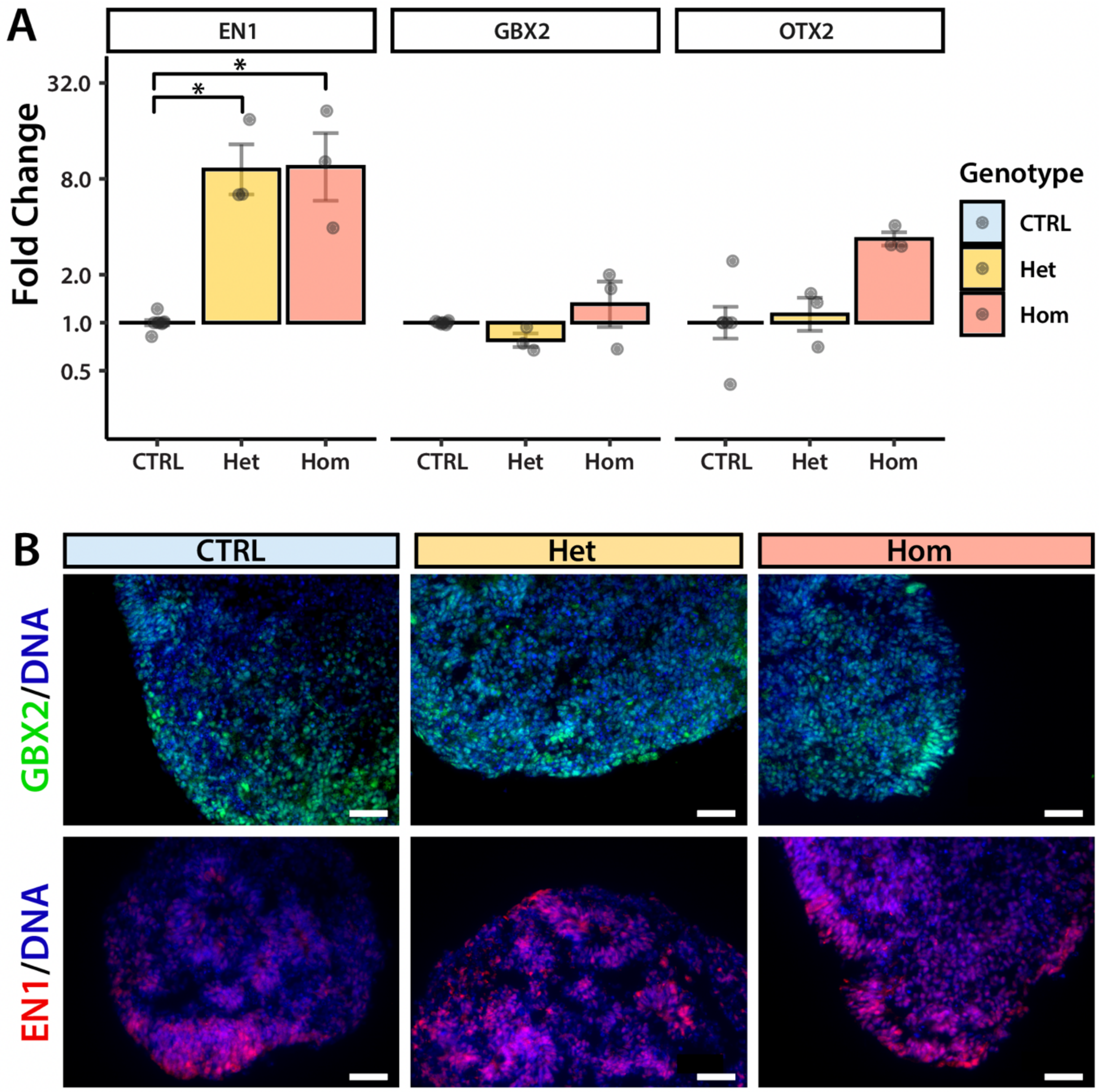

**Figure S6 Midbrain-hindbrain markers are expressed by *PTCH1* mutant organoids (related to Figure 2)**
(A) Expression of *EN1, GBX2* and *OTX2* at Day 21 relative to iPSC in heterozygous (Het) and homozygous (Hom) organoids as measured by RT-qPCR and normalised to *GAPDH* and *ACTB*. Data are composed of three or more biological replicates from separate differentiations. Statistical significance indicated as *$p<0.05$, computed by ANOVA with Dunnett's post-hoc test.
(B) Immunofluorescence staining of Day 21 organoids with antibodies specific to the hindbrain markers GBX2 (green) and EN1 (red). Nuclei are visualised in blue by Hoechst staining. Scale bars 150µm.

**Table S1 Primers used in this study**

| Gene | Fw/Rv | Sequence |
|---|---|---|
| *ACTB* | Fw | GCCGGGACCTGACTGACTAC |
| *ACTB* | Rv | TTCTCCTTAATGTCACGCACGAT |
| *ATOH1* | Fw | TGTTATCCCGTCGTTCAACAAC |
| *ATOH1* | Rv | TGGGCGTTTGTAGCAGCTC |
| *BARHL1* | Fw | TTCTGATCAGGGACATCCTTGCC |
| *BARHL1* | Rv | TGCCACTTTTGTCCAGCTTGTC |
| *CNTN2* | Fw | CCCAGCAGAGACCTATGCAC |
| *CNTN2* | Rv | ACTTTGCGCCACTTGATCC |
| *EN1* | Fw | CGTGGTCAAAACTGACTCGC |
| *EN1* | Rv | CGCTTGTCCTCCTTCTCGTT |
| *GAPDH* | Fw | GGAAGGTGAAGGTCGGAGTC |
| *GAPDH* | Rv | GTTGAGGTCAATGAAGGGGTC |
| *GBX2* | Fw | AAAGAGGGCTCGCTGCTC |
| *GBX2* | Rv | GGTCGTCTTCCACCTTTGAC |
| *GLI1* | Fw | CCCAGTACATGCTGGTGGTT |
| *GLI1* | Rv | GCTTTACTGCAGCCCTCGT |
| *KIRREL2* | Fw | CCTGAAGAAGAGGAGACAGGC |
| *KIRREL2* | Rv | TCCTCCAGAACCAGATCACTG |
| *MYCN* | Fw | GACCACAAGGCCCTCAGTACC |
| *MYCN* | Rv | TGACCACGTCGATTTCTTCCT |
| *NANOG* | Fw | GCAGAAGGCCTCAGCACCTA |
| *NANOG* | Rv | AGGTTCCCAGTCGGGTTCA |
| *NEUROD1* | Fw | GGTGCCTTGCTATTCTAAGACGC |
| *NEUROD1* | Rv | GCAAAGCGTCTGAACGAAGGAG |
| *NKX2.1* | Fw | CAAAGGCCAAACTGCTGGAC |
| *NKX2.1* | Rv | TGAGATTGGATGCGCTTGGT |
| *NKX2.2* | Fw | AGCTTCGCTTCTTTGCCTCT |
| *NKX2.2* | Rv | GGGGTCGGTCTTTTTCTCGT |
| *OCT4* | Fw | GCTCGAGAAGGATGTGGTCC |
| *OCT4* | Rv | CGTTGTGCATAGTCGCTGCT |
| *OTX2* | Fw | CACTTCGGGTATGGACTTGC |
| *OTX2* | Rv | GTGAACGTCGTCCTCTCCC |
| *PAX6* | Fw | AGCTAGCTCACAGCGGGG |
| *PAX6* | Rv | TCTGATGGAGCCAGTCTCGT |
| *PDE1C* | Fw | GAGTCGCCAACCAAGGAGAT |
| *PDE1C* | Rv | GACCGTAATCTCTGGGACGTT |
| *PTCH1* | Fw | TACTGCTCACACATCAGCCAG |
| *PTCH1* | Rv | AGTTCAGCAAGGGCAATTCAA |
| *PTCH1* | Fw | GGCAGCGGTAGTAGTGGTGTTC |
| *PTCH1* | Rv | TGTAGCGGGTATTGTCGTGTGTG |
| *PTCH1* | Fw | ACATGTACAACAGGCAGTGGA |
| *PTCH1* | Rv | ATTTCGCCCCTTCCCAGAAG |

| | | |
|---|---|---|
| *PTCH1* | Fw | TTTGCGGTGGACAAACTTCG |
| *PTCH1* | Rv | TTCAGCATTTCCTCCCAGCT |
| *PTCH2* | Fw | GATGGGGCCATCTCCACATT |
| *PTCH2* | Rv | CGCCGCAAAGAAGTACCTTACA |
| *SHH* | Fw | CTCGCTGCTGGTATGCTCG |
| *SHH* | Rv | ATCGCTCGGAGTTTCTGGAGA |
| *ZIC1* | Fw | GCCCTTCAAAGCCAAATACA |
| *ZIC1* | Rv | AGCCCTCAAACTCGCACTT |
| *ZIC3* | Fw | ATCTGCAAAGTGTGCGACAA |
| *ZIC3* | Rv | ATAGCCTGAACTGGCAGCAG |

**Table S2 Primary antibodies used**

| Target | Catalogue number | Manufacturer | Host species | Dilution |
|---|---|---|---|---|
| ATOH1 | AB5692 | Chemicon | Rabbit | 1:200 |
| EN1 | ab190080-200ul | Abcam | Rabbit | 1:250 |
| GBX2 | H00002637-M01 | Novus bio | Mouse | 1:200 |
| KIRREL2 | AF2930 | RnD systems | Goat | 1:500 |
| MKI67 | Ab16667 | Abcam | Rabbit | 1:1,000 |
| PAX6 | GTX113241 | Genetex | Rabbit | 1:1,000 |
| CCNB1 | 05-373 | Milipore | Mouse | 1:1,000 |
| TUBB3 | GT1338 | Genetex | Mouse | 1:1,000 |
| NKX2-2 | ab191077 | Abcam | Rabbit | 1:1,000 |
| CALB1 | 300 | Swant | Rabbit | 1:1,000 |

**Table S3 Secondary antibodies used**

| Target species | Host species | Conjugate | Catalogue number | Manufacturer | Dilution |
|---|---|---|---|---|---|
| Rabbit | Goat | Alexa Fluor 594 | A11037 | Invitrogen | 1:1,000 |
| Mouse | Goat | Alexa Fluor 488 | A11029 | Invitrogen | 1:1,000 |
| Goat | Donkey | Alexa Fluor 594 | A11058 | Invitrogen | 1:1,000 |
| Rabbit | Donkey | Alexa Fluor 488 | A21206 | Invitrogen | 1:1,000 |

### Supplemental Experimental Procedures

#### CRISPR clone quality control

DNA was extracted using the DNeasy kit (Qiagen, Cat#69504), and DNA was resuspended in 50µL TE buffer. After isolation, genomic DNA was subjected to PCR amplification of the target region followed by gel extraction (Monarch® Gel Extraction Kit, NEB, Cat#T1020) and Sanger sequencing. Genomic DNA from the master stocks of each clone was submitted to a genotyping array to test for gross chromosomal aberrations (Illumina Infinium Global Screening Array-24 v3.0). Each clone was tested for mycoplasma using the MycoAlert PLUS Mycoplasma detection kit (Lonza, Cat#LT07-703). Pluripotency was measured by flow cytometry using antibodies against Tra-1-60 (Biolegend, Cat#330614, 1:100), NANOG (Cell Signalling, Cat#5448S, 1:100) or isotype control antibodies (IgM 488 Biolegend, Cat#401617 or IgG 647, Cat#2985S, both 1:100).

#### EdU proliferation assay

The proliferation of *PTCH1*+/- and *PTCH1*-/- iPSC clones was determined using the Click-iT® Plus EdU Alexa Fluor® 647 Flow Cytometry Assay Kit (Invitrogen™, Cat#C10634). Proliferation was determined following the manufacturer's protocol and using 2 hours of incubation with 10µM of 5-Ethynyl-2′-deoxyuridine (EdU) for 2 hours. Proportions of cells in G0-1, S and G2-M phase were determined by co-staining with Hoechst and analysis by flow cytometry. A total of 50,000 cells per sample were analysed.
